# Ancient RNA from Late Pleistocene permafrost and historical canids shows tissue-specific transcriptome survival

**DOI:** 10.1101/546820

**Authors:** Oliver Smith, Glenn Dunshea, Mikkel-Holger S. Sinding, Sergey Fedorov, Mietje Germonpre, Hervé Bocherens, M.T.P. Gilbert

**Affiliations:** Section for Evolutionary Genomics, Department of Biology, University of Copenhagen, Copenhagen, Denmark; Greenland Institute of Natural Resources, Nuuk, Greenland; Mammoth Museum, Institute of Applied Ecology of the North of the North-Eastern Federal University, Yakutsk, Russia; Directorate Earth and History of Life, Royal Belgian Institute of Natural Science, Brussels, Belgium; Department of Geosciences, Palaeobiology, University of Tübingen, Tübingen, Germany; Senckenberg Centre for Human Evolution and Palaeoenvironment, University of Tübingen, Tübingen, Germany; Norwegian University of Science and Technology, University Museum, 7491 Trondheim, Norway

## Abstract

While sequencing ancient DNA from archaeological material is now commonplace, very few attempts to sequence ancient transcriptomes have been made, even from typically stable deposition environments such as permafrost. This is presumably due to assumptions that RNA completely degrades relatively quickly, particularly when dealing with autolytic, nuclease-rich mammalian tissues. However, given the recent successes in sequencing ancient RNA (aRNA) from various sources including plants and animals, we suspect that these assumptions may be incorrect or exaggerated. To challenge the underlying dogma, we generated shotgun RNA data from sources that might normally be dismissed for such study. Here we present aRNA data generated from two historical wolf skins, and permafrost-preserved liver tissue of a 14,300-year-old Pleistocene canid. Not only is the latter the oldest RNA ever to be sequenced, but also shows evidence of biologically relevant tissue-specificity and close similarity to equivalent data derived from modern-day control tissue. Other hallmarks of RNA-seq data such as exon-exon junction presence and high endogenous ribosomal RNA content confirms our data’s authenticity. By performing independent technical replicates using two high-throughput sequencing platforms, we show not only that aRNA can survive for extended periods in mammalian tissues, but also that it has potential for tissue identification, and possibly further uses such as *in vivo* genome activity and adaptation, when sequenced using this technology.

## Introduction

The recent revolution in the sequencing of ancient biomolecules has allowed multiple layers of -omic information – including genomic [1], epigenomic [2, 3], metagenomic [4, 5], and proteomic [6, 7] – can be gleaned from ancient and archaeological material. This raft of evolutionary information almost all derives from either DNA or protein, biomolecules both traditionally thought to be considerably more stable than RNA. This is unfortunate, since transcriptome data has the potential to access deeper layers of information than genome sequencing alone. Most notably these include assessments of the *in vivo* activity of the genome, and assessing other aspects of ancient bio-assemblages such as biotic colonisation / microbiomes [8], host-pathogen interactions [9], and the level of post-mortem molecular movement within remains and surrounding media [10].

Despite the dominance of DNA, in recent years several studies have begun to explore whether or not RNA survives in archaeological substrates, particularly in the context of plant materials. Next-generation sequencing (NGS) approaches have uncovered viral RNA genomes in barley grains and fecal matter [11, 12], environmentally-induced differential regulation patterns of microRNA and RNA-induced genome modifications in barley grain [13, 14], and general transcriptomics in maize kernels [15]. All but one of these datasets however has been derived from plant seed endosperm, which often facilitates exceptional preservation [16, 17] and is known to be predisposed to nucleic acid compartmentalisation [18], thus allowing for reasonable expectations of such preservation. The conjecture that ribonucleases released during soft tissue autolysis would virtually annihilate RNA had, until recently, discouraged researchers from attempting such sequencing in animal tissues in favour of more stable molecules. This is exemplified by the fact that to date, ancient RNA data has been generated directly from ancient animal (human) soft tissues in only one example [19], and this was without utilising NGS technology. Instead, a targeted qPCR approach was used, presumably intended to bypass extraneous noise that might be expected in ancient NGS datasets. The recent qPCR-based approach to microRNA identification demonstrated persisting specificity in permafrost-preserved human tissues [19] and thus opened the possibility of a more complete reconstruction of ancient transcripts in soft tissues when preserved under favourable conditions. While complexities surrounding the survival of purified RNA within a long-term laboratory storage setting are well documented [20, 21], the complex thermodynamics of RNA lability and enzymatic interactions are themselves not well understood, especially within long-term post-mortem diagenesis scenarios [22]. Evidence exists that suggests that the survival of purified (modern) RNA is influenced by the specific tissue from where it originated [23], suggesting co-extraction of tissue-specific RNases is a significant problem. Others have suggested that the chemical structure of RNA is such that its theoretical propensity for spontaneous depurination is less than that of DNA [24]. Although strand breakage should occur more often, the observable depletion of purified RNA within a laboratory setting has often been attributed to contamination from RNases which are often active in purified samples even when frozen. Because chemical and enzymatic interactions in archaeological or paleontological assemblages are generally unpredictable at the molecular level, it is possible that the activity of RNAses, and the susceptibility of RNA to those enzymes within a complex matrix of biomatter, could be slowed or arrested through uncharacterised chemical interactions. As such, it is possible that under environmental conditions such as desiccation or permafrost, ancient RNA may indeed persist over millennia. Exceptionally-well preserved remains provide an opportunity to test this hypothesis. Given this, we decided to take advantage of some recently recovered samples exhibiting a range of ages and DNA preservation [25]. We felt these were ideal animal candidates to test for both the persistence of ancient RNA in such contexts. The results presented here describe the oldest directly sequenced RNA, by a significant margin, alongside younger tissues which still may be seen as novel substrates given the prevailing RNA dogma. To confirm the absence of platform-specific biases, we sequenced each sample using the Illumina HiSeq-2500 platform and performed a technical replicate (library and sequencing) on the BGISEQ-500 platform. For clarity, the biological results and interpretations shown in the main text refer to HiSeq-2500 data since Illumina sequencing platforms are the most often used for sequencing ancient DNA, with BGISEQ-500comparisons referenced directly where necessary and in the supplementary materials. From the results presented here, we propose that the range of aRNA sources now extends to both animals and plants, thus opening up the possibility of routinely using ancient RNA as a valuable biomolecular resource for future research.

## Results

### RNA recovery and sequence data from ancient tissues

From between 47mg and 665mg of tissues including skin, cartilage, liver, and skeletal muscle, we recovered between 100ng and 461ng RNA (see Table 1). Unsurprisingly, there was a marked difference between the ancient Tumat and historical samples: while the historical skin samples gave between 3.4µg and 6.7µg RNA per gram tissue, the ancient Tumat samples only gave between 0.28µg and 0.57µg per gram. After sequencing and mapping, we calculated the endogenous RNA content of the tissues to be between 7.4% - 80.0% using the HiSeq-2500 platform (Table 2).

**Table 1:**
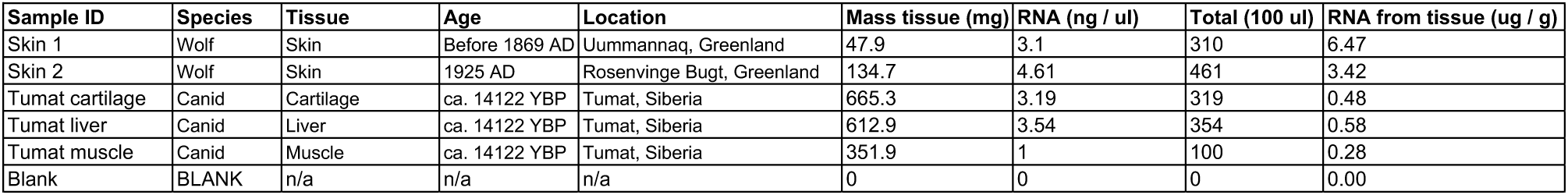
Basic sample details including age, tissue, and RNA extraction statistics.

| Sample ID | Species | Tissue | Age | Location | Mass tissue (mg) | RNA (ng / ul) | Total (100 ul) | RNA from tissue (ug / g) |
| --- | --- | --- | --- | --- | --- | --- | --- | --- |
| Skin 1 | Wolf | Skin | Before 1869 AD | Uummannaq, Greenland | 47.9 | 3.1 | 310 | 6.47 |
| Skin 2 | Wolf | Skin | 1925 AD | Rosenvinge Bugt, Greenland | 134.7 | 4.61 | 461 | 3.42 |
| Tumat cartilage | Canid | Cartilage | ca. 14122 YBP | Tumat, Siberia | 665.3 | 3.19 | 319 | 0.48 |
| Tumat liver | Canid | Liver | ca. 14122 YBP | Tumat, Siberia | 612.9 | 3.54 | 354 | 0.58 |
| Tumat muscle | Canid | Muscle | ca. 14122 YBP | Tumat, Siberia | 351.9 | 1 | 100 | 0.28 |
| Blank | BLANK | n/a | n/a | n/a | 0 | 0 | 0 | 0.00 |

**Table 2:** NGS data and mapping summary, with calculations of endogenous content and RNA enrichment factors.

|  | Sample # | Species | Tissue | Age | Total reads post-adaptor trimming | Genome | mRNA | rRNA | Proportion rRNA | tRNA | RNA Enrichment factor | Endogenous % |
| --- | --- | --- | --- | --- | --- | --- | --- | --- | --- | --- | --- | --- |
| BGISEQ | Skin 1 | Wolf | Skin | Before 1869 AD | 69,053,233 | 26,043,866 | 6,858,947 | 16,714,271 | 31.03% | 4,243,690 | 14.69 | 37.72% |
|  | Skin 2 | Wolf | Skin | 1925 AD | 6,675,338 | 5,581,322 | 1,288,462 | 4,696,537 | 39.40% | 354,381 | 15.62 | 83.61% |
|  | Tumat C | Canid | Cartilage | ca. 14122 YBP | 44,765,013 | 2,244,289 | 783,522 | 401,982 | 11.61% | 32,077 | 7.46 | 5.01% |
|  | Tumat L | Canid | Liver | ca. 14122 YBP | 27,626,403 | 16,509,691 | 5,038,336 | 3,570,007 | 10.91% | 7,617,698 | 13.52 | 59.76% |
|  | Tumat M | Canid | Muscle | ca. 14122 YBP | 66,780,343 | 3,815,483 | 1,057,959 | 1,357,348 | 20.73% | 317,792 | 9.85 | 5.71% |
|  | Blank | BLANK | n/a | n/a | 1,701,272 | 56,822 | 20,808 | 126,467 | 55.43% | 24,069 | 41.47 | 3.34% |
| HiSeq | Skin 1 | Wolf | Skin | Before 1869 AD | 23,258,645 | 11,366,481 | 3,493,902 | 7,612,932 | 31.83% | 1,441,633 | 15.18 | 48.87% |
|  | Skin 2 | Wolf | Skin | 1925 AD | 32,927,602 | 26,320,301 | 5,618,346 | 19,883,788 | 36.95% | 1,990,974 | 14.36 | 79.93% |
|  | Tumat C | Canid | Cartilage | ca. 14122 YBP | 20,915,948 | 2,354,199 | 1,064,732 | 209,067 | 5.71% | 31,676 | 7.63 | 11.26% |
|  | Tumat L | Canid | Liver | ca. 14122 YBP | 6,811,527 | 4,114,476 | 1,882,220 | 1,192,800 | 14.94% | 796,571 | 12.94 | 60.40% |
|  | Tumat M | Canid | Muscle | ca. 14122 YBP | 39,878,232 | 2,932,798 | 1,099,000 | 818,537 | 16.44% | 127,563 | 9.59 | 7.35% |
|  | Blank | BLANK | n/a | n/a | 1,339,288 | 75,612 | 91,929 | 9,498 | 5.33% | 1,029 | 18.63 | 5.65% |

### RNA enrichment

For each sample, we took frequencies of individual reads mapping to the entire genome, and similarly the frequencies of individual reads mapping to only the transcribed regions of the genome (mRNA, rRNA and tRNA). We then divided the RNA read frequency with the whole-genome read frequency for each sample to give an enrichment factor (Table 2). We found between 7.4-fold and 15.6-fold enrichment for transcripts from HiSeq-2500 data. We found no significant age- or tissue-related correlation to enrichment level.

We subjected earlier DNA sequencing data from the same samples used in this paper [25] to the same transcriptome mapping pipeline as our RNA data, in order to confirm that the enrichment of transcriptomic reads we saw in the RNA data was not spurious or the result of DNA contamination. As with the RNA data, we calculated the RNA enrichment factor for each sample. Whereas we saw at least 7.4-fold transcript enrichment for the RNA data, we saw only between 0.2- and 1.2-fold enrichment for the equivalent DNA data. Further, while the RNA data showed that a large proportion of the endogenous content for each sample (between 5.7% and 37%) was of ribosomal origin, the ribosomal content of the endogenous DNA was significantly lower, between 0.09% and 0.15%, and we suspect more likely a representation of rRNA genes than their transcripts. Considering this, and the known high abundance of rRNA as a proportion of cellular RNA, this strongly suggests that the RNA-seq dataset represents authentic RNA, with minimal, if any, DNA contamination.

### Junction analysis

To further establish that we had sequenced RNA, as opposed to contaminant single-stranded DNA (ssDNA), we assessed the frequencies of reads straddling intron-exon (splice) junctions and those straddling exon-exon junctions. With RNA-seq data, we would expect to observe a high proportion of exon-exon reads to demonstrate that precursor mRNA processing has taken place in active transcripts, but we would also expect to see a degree of intron/exon reads representing precursor mRNA themselves. We found that in all cases, the number of reads mapping to exon/exon junctions was greater, often by orders of magnitude, than those mapping to splice junctions (Table S1). In particular, the Skin #2 and Tumat liver samples respectively showed 186-fold and 68.5-fold more reads mapping to exon-exon junctions than splice junctions. We then repeated this analysis using DNA data generated from the same samples, as a negative control [25]. We found the DNA data showed the opposite trend to RNA-seq data, with exon-exon junctions being significantly under-represented compared to splice junctions in all cases. These analyses further suggest that our primary data represents authentic aRNA.

### Damage profiles

Damage profiles were not consistent with typical ancient DNA profiles, although the expectations for comparing RNA and DNA in this manner are unknown due to a general lack of aRNA NGS data. mapDamage analysis of earlier DNA sequencing of the same samples showed profiles that were typical of ancient DNA, although at low levels for samples as old as the Tumat canid. Unsurprisingly, the two samples with the lowest levels of damage were the historical skin tissues. Interestingly, the liver sample, which showed the greatest affinity to its modern counterpart in transcriptome analysis, had the lowest damage levels of all tissues from the Tumat canid, further suggesting its exceptional preservation.

The RNA profiles themselves showed either low-levels of damage throughout when de-duplicated, and some elevated C > U transitions towards the centre of the molecule (supplementary Figure S1). Interestingly, the damage appears at lower levels than the equivalent DNA samples. The damage was generally limited to C > U misincorporations as opposed to G > A misincorporations, which is consistent with data deriving from a single-stranded library construct. Damage patterns were more pronounced when duplicates were retained, which is unsurprising considering the level of sequence duplication. We also note that the damage in general is more pronounced in data from the HiSeq-2500 platform.

### Statistical inter- and intra-tissue comparisons of ancient transcriptomes (method 1)

Over the entire dataset ordination and clustering revealed that the ancient samples were globally more similar to each other than to the control samples and vice versa (Supplementary Figures S2 and S3). However, when considering individual ancient / historical samples against all control samples, we found that the ancient Tumat liver and historical Skin 2 samples were most similar to their modern counterparts. Clustering also revealed a set of 71 genes with relatively highly abundant transcripts across all, or most ancient samples in comparison to the control samples (Supplementary Table 2).

Considering the most highly expressed genes (i.e. 95^th^ percentile) in each control tissue, there were some relationships of note between control and ancient samples. There was a significant relationship between control liver and ancient liver, with control liver expression explaining 16% (Adjusted R^2^ values) of the variation in ancient liver transcript abundance (Supplementary Data S1; Figure 1). Control liver gene expression was more similar to ancient liver transcript abundance in comparison to any of the other ancient samples or any of the other control samples (Supplementary Data S1). Similarly, there was a significant relationship between control skin gene expression and transcript abundance in the historical Skin 2 sample, with control skin expression explaining 8% of the variation in historical Skin 2 transcript abundance (Supplementary Data S1; Supplementary Figure 4). There was also a marginally significant relationship between control skin and historical Skin 1 (P = 0.012, α = 0.01), however it explained only a very small amount of the variation in Skin 1 transcript abundance (0.4%; Supplementary Data S1). Control skin gene expression was more similar to both historical skin sample transcript abundance(s) in comparison to any of the other ancient samples, however there were also significant relationships with all other control tissues (Supplementary Data S1). There was no relationship between control cartilage gene expression and ancient cartilage transcript abundance, although there was a relationship with Skin 2 transcript abundance, control liver and control skin gene expression (Supplementary Data S1). There were no significant relationships between control muscle gene expression and any of the ancient samples or the other control samples. All pairwise regression parameters and details are provided in Supplementary Data S1.

**Figure 1:**
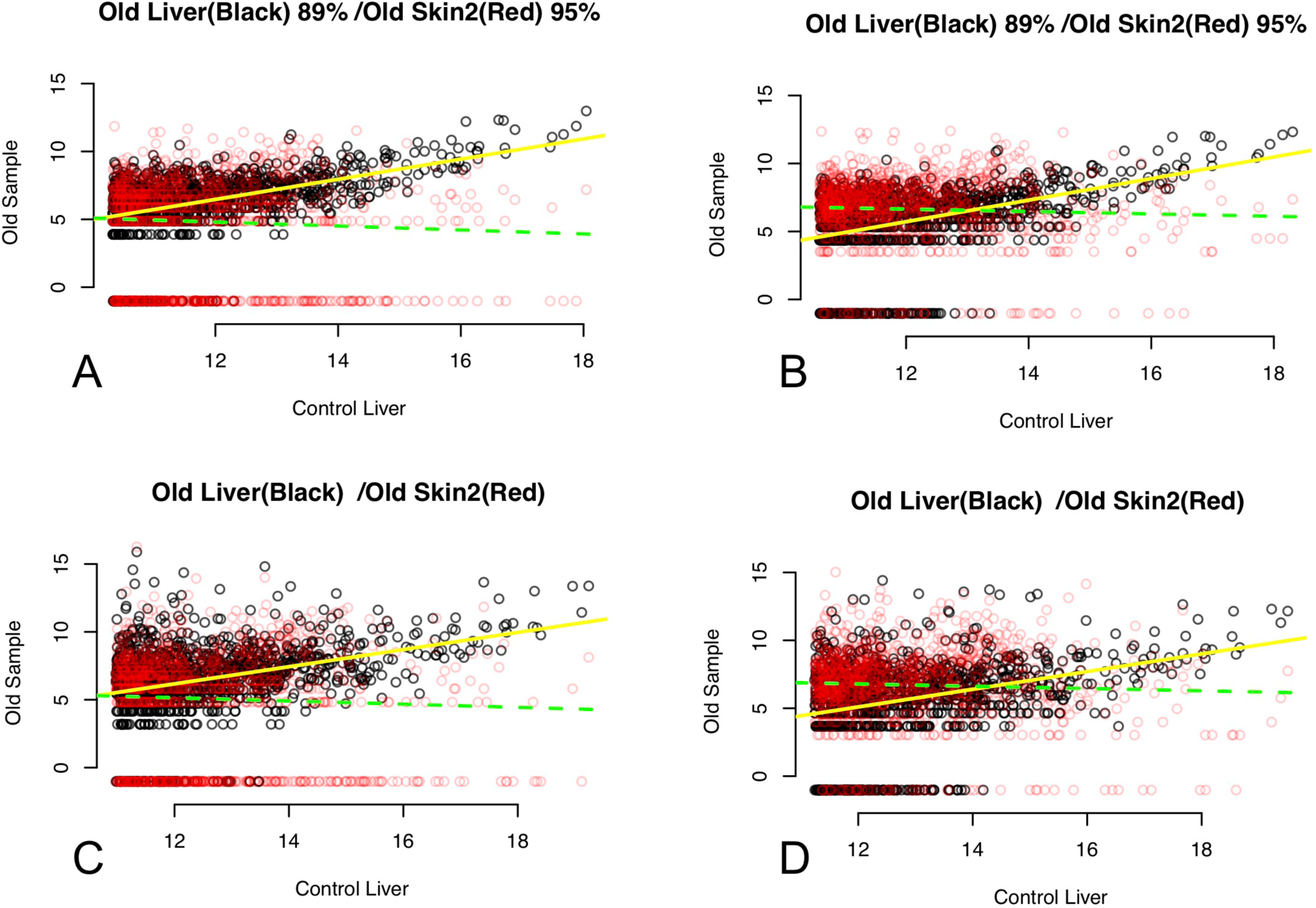
Regressions of ancient liver and historical skin samples, method 1: relationships between 95% percentile of expressed genes in each control tissue sample (x-axis, graph title) and each ancient sample or control samples from other tissues (y-axis, graph title). Black points in graphs comparing ancient samples are the relationship between the control tissue and the same ancient tissue, red points overlaid are the relationship between the control tissue and other ancient tissues (in graph title – one per graph). Yellow lines are least squares linear regression fit for black points and green lines are least squares linear regression fit for red points. Filled lines indicate a significant linear regression, dashed lines indicate a non-significant linear regression. A) BGISEQ-500 data, de-duplicated; B) HiSeq-2500 data, de-duplicated; C) BGISEQ-500 data, duplicates retained; D) HiSeq-2500 data, duplicates retained.

### Tissue specificity when compared to the Canine Normal Tissue Database (method 2)

Like our observations from Method 1, we found that the historical Skin 2 and the ancient Tumat liver tissues showed significantly more similarity to their modern control counterparts than the other historical / ancient tissues. Of the 14,300 years old Tumat samples, we found virtually no correlation between ancient and control data when compared to the canine normal tissue array (method 2) using muscle (r^2^ = 0.07) and cartilage (r^2^ = 0.01). However, we observed a high degree of similarity with liver tissue, when similarly compared to modern data (r^2^ = 0.94, Figure 3). We immediately noted that several highly-expressed genes in the ancient liver tissue are associated with liver function including apolipoproteins, fetuins, and retinol-binding proteins.

**Figure 2:**
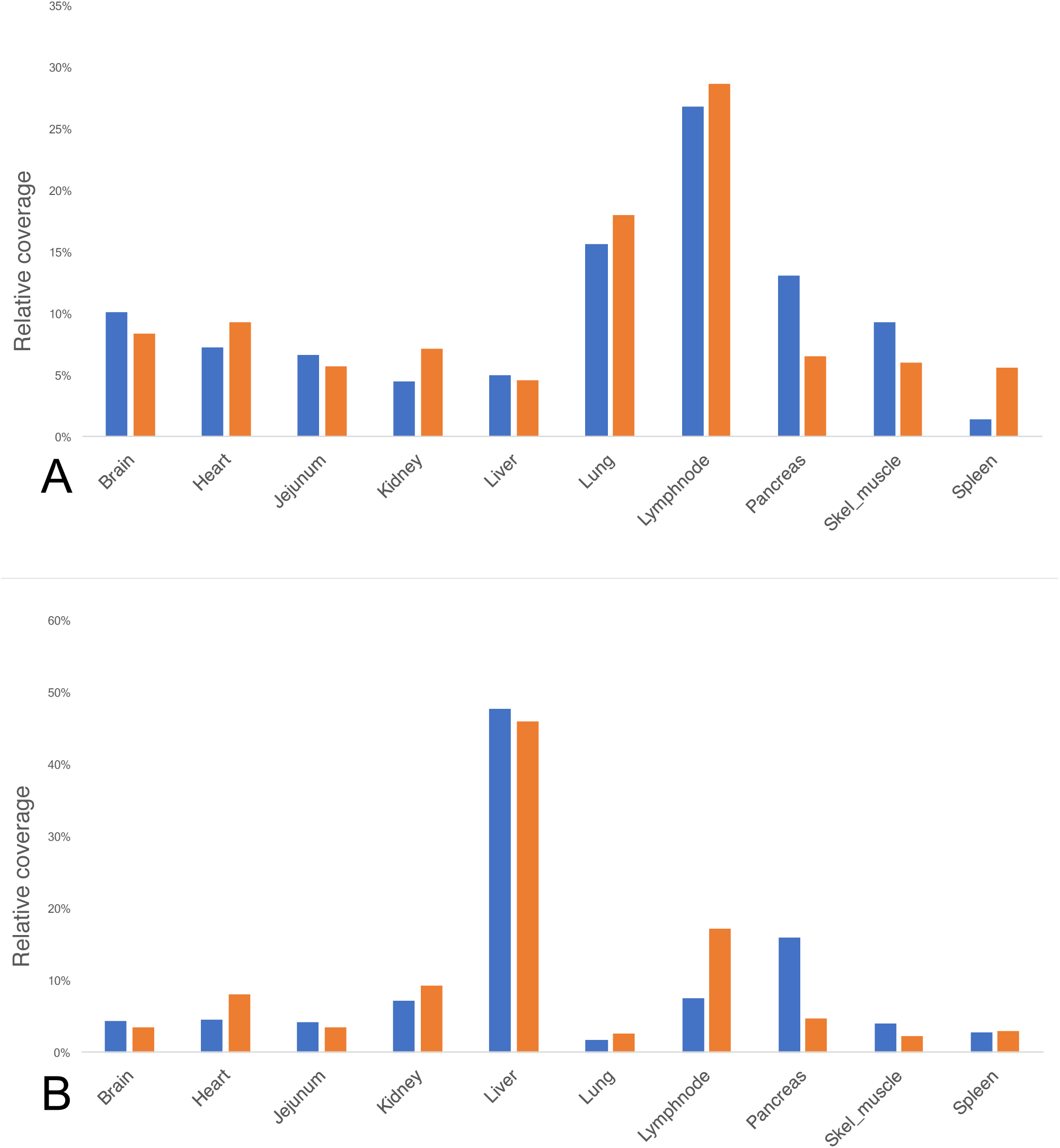
Comparison of ancient and control tissues using Method 2. Coverage scores (Y-axis) were calculated based on the mean coverage of reads to each named gene in the CanFam3.1 transcriptome, followed by filtering to the 95^th^ percentile of all genes represented. Each gene was then assigned a most-associated tissue based on data Affymetrix array derived from 10 canine tissues (X-axis). Each tissue was then assigned a cumulative score based on the coverage scores of each gene in the 95^th^ percentile. Orange bars represent modern control tissues and blue bars represent ancient / historical tissues. Panel A: historical Skin 2 versus control skin. Panel B: ancient Tumat liver versus control liver.

**Figure 3:**
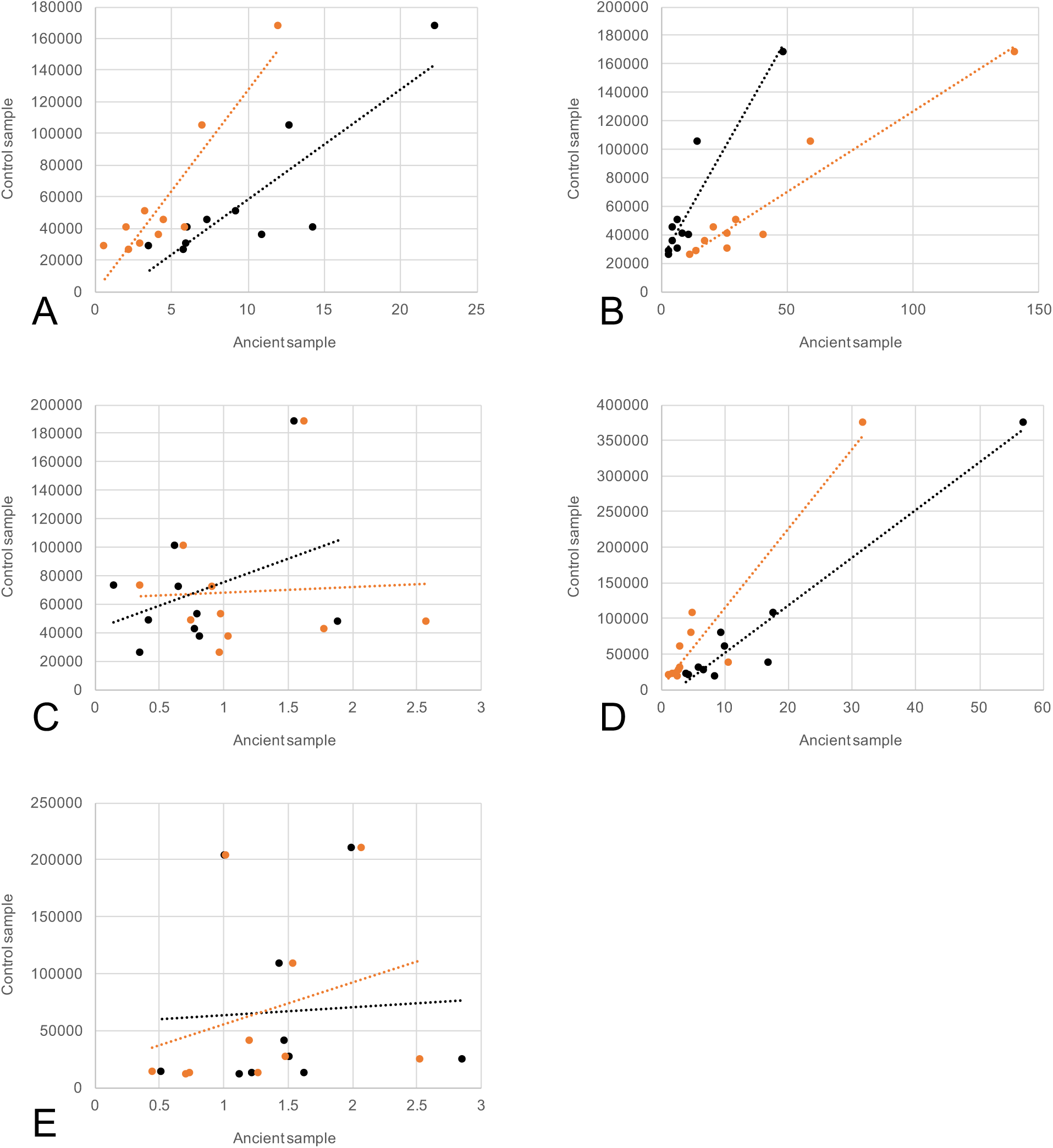
Regressions of all samples, method 2: Relationships between 95% percentile of expressed genes in ancient tissues (x-axis) versus control samples (y-axis). Values are calculated based per-tissue scores (see methods) having removed duplicate reads from mapping data. Black data points and trendline refer to BGISEQ-500 data, while orange data points and trendline refer to Illumina HiSeq-2500 data. A) Skin 1; B) Skin 2; C) Tumat cartilage; D) Tumat liver; E) Tumat muscle

A high level of similarity between historical and modern skin tissues (r^2^ = 0.70 for Skin 1 and 0.87 for Skin 2) was also observed using method 2 (Figure 3). We noted that highly-expressed genes in both ancient and controls are associated with skin and connective tissue, including collagen and several keratin-producing genes (supplementary Table S2).

### GC content and read duplication

The GC content of full reference transcripts falling within the 95^th^ percentile of abundance was between 51% and 57% (Supplementary Table S3). We noted that the GC content of reads mapping to those transcript sets exhibited higher GC content than the transcripts themselves, which is not unexpected considering previous aRNA results [13, 15, 19]. On average, the de-duplicated datasets had 4.6% greater GC content than the references, and the redundant (i.e. duplicates retained) datasets showed on average 7.3% higher GC content. This suggests a slight bias towards high-GC fragments being preserved, which is again not unexpected in RNA-seq data, given that transcribed regions of the genome are generally GC-rich [26]. However, the uniquely short nature of read fragments, compared to a modern RNA dataset, combined with non-uniform GC content across a given transcript, suggests that the GC bias observed here does not skew the resulting transcription profiles.

Due to the high number of PCR cycles (20) required to build libraries, it is unsurprising that we observed significant duplicate reads in all ancient samples, between 80.9% and 87.1%. However at least some of this variance can be explained by ‘true’ transcript abundance, exemplified by the control data from modern material being between 20.9% and 39.4% duplicate reads.

### Metagenomic analysis

To explore microorganism presence, and further validate the authenticity of our RNA reads, we performed two metagenomic analyses. First, on the tRNA fraction, to validate the origin of the data as being canine due to the relatively high interspecies sequence divergence of tRNA; we found that in all cases, the vast majority (> 86.5%) of reads were assigned either directly or directly basal to canine tRNA, further suggesting the authenticity of our data.

Secondly, we looked for evidence of viral infection from RNA viruses (both ssRNA and dsRNA) in all the sequenced tissues, noting that previous aRNA work has revealed RNA viral genomes in ancient material [11, 27]. We found no evidence of viral sequences in our RNA data.

## Discussion

Our results show the proof-of-principle that under permafrost conditions, tissue-specific transcriptome profiles are potentially recoverable from mammalian soft tissues preserved over thousands of years. Since the survival of RNA for such long periods of time is unexpected, verification of the data’s authenticity is important. By comparing the RNA data to equivalent DNA data and assessing key characteristic differences between RNA and DNA data such as reads mapping splice junctions versus exon-exon junctions, the quantity of ribosomal RNA in the samples, and overall transcriptome enrichment, we have shown the expected differences to be present and thus believe the data presented here is truly representative of ancient RNA.

We suggest that in contexts conducive to biomolecular preservation, ancient RNA (or ‘palaeotranscriptome’) analysis could provide a number of standard additional facets to the biomolecular archaeological toolkit. With further research, we anticipate these could be expanded to include tissue identification, metagenomic palaeopathology of RNA viruses, and identifying specific *in vivo* processes concerned with individual genomes and their underlying causes, such as climate, diet, trauma, and disease.

### Tissue specificity in ancient tissues

Of the 2 historical skin samples and 3 ancient tissue samples, 2 samples (Skin #2 and Tumat liver) exhibited signals strongly associated with their modern counterparts. The ancient liver sample in particular, despite being the oldest of the three individuals, showed the greatest similarity to its control sample. Of particular note is that when compared to the reference Affymetrix array using method 2, prior to comparative analysis with the control sample, 80% of the 10 most abundant transcripts and 50% of the 50 most abundant transcripts are biologically sensible, i.e. are genes primarily associated with liver tissue. Within those 50, 5 were class A and C apolipoprotein isoforms involved in lipid transport and, crucially, synthesised within the liver [28]. Three different isoforms of alpha-2 glycoprotein, associated with liver function in mammals [29] were present, as were several fibrinogen and fetuin-B genes which are also liver-derived [30, 31]. While simple identifications such as these are by no means conclusive, we took them as a starting point to perform more detailed statistical analyses. However, we noted that far from being an isolated incident, other, different tissues exhibited similar superficial equivalence to their controls. The skin 2 sample contained 19 keratin-associated isoforms within the most abundant 50 transcripts, alongside several proline-rich proteins, both of which are associated with dermal tissue. Several microRNA genes were also highly represented, although a reference set for canine microRNA tissue-specificity does not include skin [32] and so concrete conclusions about those transcripts cannot be made.

In addition to tissue differentiation, it was encouraging to note that in all tissues, the most highly-expressed gene without tissue-specific assignment in our scoring matrix was the RN7SL1 cytoplasmic RNA, which forms part of the ribosomal nucleosome complex. In highly degraded tissues, the significant presence of ribosomal RNA (rRNA) is expected [15] and therefore is further evidence of RNA enrichment. Ribosomal RNA (rRNA) itself accounted for between 5.7% and 39.4% of the reads, again with no obvious correlation to tissue type or age, but again with similar results between sequencing platforms (r^2^ = 0.90). Similarly, all ostensibly connective tissues included a predicted collagen alpha-like gene (LOC102152155) as the second- or third-most expressed locus, although a specific named homologue could not be identified for downstream statistical analysis.

### Ancient RNA preservation in permafrost and historical tissues

While the sample set is small, we noted that the ostensibly best-preserved tissue in the Tumat #2 individual is the deepest (liver), and the least well-preserved is the most superficial (cartilage). The muscle tissue, while intermediate, was closer in quality to the cartilage. Although we cannot make a confident assertion, we suspect that, at least concerning a small animal preserved in permafrost, the deepest tissues might have a higher proportion of endogenous DNA / RNA because of the fact that external microbial or other environmental activity would be initially present on the outer tissues. This is reflected in the lesser endogenous content of the outer tissues. Microbial activity on surface tissues being arrested by rapid freezing before reaching deeper tissues would also explain the higher endogenous content of the liver. It is also logical that a transcriptionally active tissue such as liver would exhibit greater specificity through time due to the absolute (as opposed to proportional) levels of nucleic acids in the tissue itself. We hypothesise that degradative enzymes in liver tissue would have no effect on the proportion of endogenous RNA given the overall rapid freezing of the animal as discussed above. With regards to historical samples, it is unsurprising that the older of the two skin tissues shows weaker RNA preservation, although this may have been affected by hitherto unknown and different preservation methods and individual post-mortem histories.

As with any extraordinary claim, the veracity of our results is hugely important. Therefore we analysed our RNA-seq data in conjunction with equivalent DNA data to eliminate the possibility of DNA contamination, by looking at exon-exon junctions, overall mapping proportions, biologically-relevant tissue-specific transcriptome activity, and ribosomal RNA content. The results of these analyses all show compelling evidence of the authenticity of the RNA data, reinforcing once more the exceptional character of these remains for palaeobiological and palaeophysiological research on extinct mammals or ancient representatives of still extant species.

### RNA damage profiles

RNA Damage profiles, while generally low-level and consistent with the equivalent DNA damage profiles (Figure S9), are less consistent with earlier observations of ancient RNA damage which show consistent high-level damage across reads with elevated C>U misincorporations at both ends [11]. However, the equivalent DNA profiles are likely to be a better proxy on which to compare these damage profiles, because the source of the other RNA (in this case, desiccated seeds from southern Egypt) is wildly different in terms of tissue (plant seed endosperm) and burial context (extreme changes in temperature including highs in excess of 40°C). Additionally, these data are some of the only available NGS data derived from aRNA available. The earlier model proposed that RNAs propensity to form secondary structure by self-folding protects mid-sequence cytosines from hydrolytic attack, whereas terminal bases are more exposed and thus more likely to become deaminated. This characteristic is also seen in single-stranded ancient DNA libraries [33], and the different profiles seen in the RNA data suggest that there is little or no DNA contamination in the canine RNA libraries. This being said, we stress that because NGS data derived from aRNA are generally rare, there are very few expectations as to what a ‘typical’ aRNA damage profile would look like. Previous transcriptome data from ancient maize kernels shows consistent, low-level damage across the strand, similar to that observed in the historical skin samples shown here [15] although less pronounced than our Pleistocene canid data. We postulate that secondary structure formation, while routinely thermodynamically predictable as *in-situ* transcripts [34], could result in inconsistent or unpredictable (dynamic) de- or re-exposure of cytosine molecules during RNA diagenesis and would thus be, unsurprisingly, a time-dependent diagenetic process. This may be compounded by stochastic fragmentation of RNA molecules, resulting in re-folding or the creation of RNA pseudoknots, the structures of which are less predictable [35]. Further data from a range of sources is needed to crystallise these expectations, and develop models to more accurately predict secondary structure formation in diagenetic assemblages.

### Sequence duplication in ancient RNA-seq data

The question of whether to de-duplicate RNA-seq data is much debated [36]; potential issues surrounding type I and type II errors, the effect of greater or fewer PCR cycles, and difficulties in distinguishing a transcript duplicate from a PCR duplicate all contribute to a general uncertainty. In practice, the prevailing opinion appears to be that decisions should be based on individual samples. Some recent developments however suggest that distinguishing duplicate types may be viable under certain circumstances, either computationally [37], or through a molecular-indexing approach [38]. The data presented here however is unique in its age and origin, generated from small starting amounts of RNA and thus prone to type I errors introduced during PCR. On the other hand, random survival of short sequences over long time periods, the effect of secondary structure formation, and other biological processes *may* lend themselves to type II errors. On balance however, we decided that the most parsimonious approach, considering the high numbers of PCR cycles required and the shorter than usual nature of the fragments, would be to treat the de-duplicated dataset as the most informative.

### GC content of ancient RNA data

We noted that the GC content of reads was slightly higher than those of the transcripts to which they were mapped, and further increased when accounting for duplicate reads (Figure S5). We believe that a combination of excess duplicates arising from the high number of PCR cycles necessary for NGS library construction (as opposed to ‘true’ transcript duplicates), the trend of transcribed regions of mammalian genomes being generally GC-rich [26] and the greater survivability of GC-rich fragments of ancient biomolecules, is responsible for this observation. We therefore suggest that in this instance, the de-duplicated datasets are more likely to be accurate approximations of the ‘true’ transcripts from these samples. We observed in both our statistical methods applied to read coverage that the de-duplicated ancient datasets showed significantly greater similarity to control dataset, regardless of de-duplication of the controls. This is likely due to the fact that duplicates in the control samples were significantly lower, and where present, representative of actual *in vivo* transcript expression as opposed to PCR biases. In all cases, the GC content was elevated in datasets with duplicates retained; however the BGISEQ-500data showed that this trend was slightly less pronounced, despite library protocols being identical apart from the platform-specific adapters used and the sequencing platform itself.

### Comparison of Illumina HiSeq-2500 and BGISEQ-500 sequencing platforms

Following the comparison of Illumina and BGISEQ-500platforms on aDNA, which showed little difference in standard quantitative metrics between them [25], we decided to use both platforms in this study to a) compare the two when using aRNA instead of aDNA, and b) treat one as a technical replicate for proof-of-concept purposes. Overall, we found very little difference between platforms in terms of sequence quality, GC bias and overall analytical outcomes between HiSeq-2500 and BGISEQ-500platforms (Figure S7), in keeping with previous comparisons of these platforms using DNA data [25]. The most noticeable difference was the fragment size distribution after adapter removal; we noted that the HiSeq-2500 gives a higher proportion of small fragments than BGISEQ-500 (Figure S8), likely due to preferential clustering of small fragments as noted previously by Illumina. Crucially however, we noted that comparisons following biologically meaningful analyses retained strong correlation. In particular, we found that the calculated endogenous content and RNA enrichment factors were almost identical for both following linear regression (r^2^ = 0.98 and 0.96 respectively, Figure S7 panels A and D, Table 2). The relationships between control and ancient tissues using Method 1 were also very similar, with BGISEQ-500slightly outperforming HiSeq-2500 explaining 20% of the variance (compared with 16% explained with HiSeq). The standardised individual gene expression metrics and similarity between individual samples were likewise similar between the two platforms (Figure S2).

In terms of GC content of mapped reads, we did note slightly higher discrepancies between the two sequencing platforms: Of the reads mapping to transcripts in the top 95^th^ percentile of coverage depth, we found lesser but significant correlation (r^2^ = 0.78), and GC of all reads following duplicate removal at a similar correlation (r^2^ = 0.75). A better correlation was observed in GC content of all reads prior to duplicate removal (r^2^ = 0.85), suggesting that both platforms gave data slightly biased towards GC retention. This is not to say the platforms themselves exhibit bias, but is more likely to be a function of long-term preservation favouring GC-rich molecules as previously noted [39]. We did however notice this bias to be slightly increased overall in the BGISEQ-500 platform (Figure S5, Figure S7 panel C), although this effect appears to be negligible in downstream analysis. We also note that the recommended library input requirements into pre-sequencing treatment are higher for BGISEQ, which is not an insignificant point considering the generally much smaller quantities of DNA / RNA available to palaeogenomic study.

In terms of read duplication, we found that the BGISEQ-500 platform slightly outperformed HiSeq-2500 by having a lower proportion of duplicated reads in all samples except Tumat liver. However, we noted that while higher, duplication levels from the HiSeq-2500 platform were more consistent with each other, varying between samples by 6.2% versus the BGISEQ-500 platform at 20.1%. Since our primary analyses and conclusions are based on de-duplicated reads, this result makes no difference to our conclusions. For the analysis of reads straddling splice or exon-exon junctions, we again found little difference between platforms, although again the BGISEQ-500 slightly outperformed HiSeq-2500 in identifying a higher proportion of exon-exon junction reads compared to splice junction reads in the RNA data. The relative proportions of the same analysis performed on the previously-sequenced DNA data showed negligible differences between the two platforms (Table S1). While both platforms are broadly similar in terms of all metrics of the data returned, we suggest that researchers, particularly those working with low-yield ancient samples, should consider issues such as data output, cost-per-read, and library input mass, to decide on the best fit for individual projects.

### The future of ancient RNA

Research using ancient biomolecules is moving in leaps and bounds, breaking barriers particularly in terms of throughput, sample age, starting material, and the range of biomolecules at our disposal. Ancient RNA, although touched upon in very recent literature, is still relatively unstudied. Perceptions about what aRNA can inform us about, that DNA or proteins cannot, and a more general instability, lead many to dismiss it as unlikely and unnecessary. These data represent the oldest ancient RNA from any source to be sequenced, by a significant margin, and show that under a range of conditions, aRNA can remain intact well enough to identify specific transcriptomic profiles approximately 9,000 years earlier than the current oldest sequenced aRNA. Previous research in plants has identified the potential to uncover aRNA viruses, and monitor *in vivo* activity in long-dead organisms, although these were exceptionally well preserved and not prone to typical enzymatic or autolytic process that occur in mammalian decomposition. This research confirms that these processes are sufficiently arrested in permafrost animal remains, and as such, *in vivo* processes can now be identified in samples of great interest to current research themes. This potential need not be limited to permafrost samples, but extending to other low-temperature climates such as Greenland, Alaska, Canada and Antarctica. Equally, source material need not be limited to soft tissues; as previous research has shown, a variety of organic materials are potential sources of aRNA (most notably seed endosperm) and so there is potential to explore aRNA preservation in bone, keratin, or even sediments from such environs. Further, we anticipate that other biomolecular analysis may be used to complement and cement our understanding of *in vivo* processes; for example, quantitative palaeoproteomic approaches, still in their infancy, could be enhanced using relative transcriptome data. Additionally, stable isotope data could further be complemented by these data; nitrogen isotopic analysis of different tissues indicate that Tumat puppy#2 was still sucking its mother’s milk when it died, and so it may be possible, with more samples, to establish individual developmental stage through transcriptomic and isotopic complementary data.

In conclusion, we suggest that as an untapped biomolecular resource, ancient RNA has great potential in enrich the current body of palaeogenomic study. Not only has it the potential to provide verification for tissue identification, but also to enhance or validate other areas of biomolecular archaeological research such as epigenomics, palaeoproteomics, and stable isotope analysis. Continuing the palaeopathological perspective, we note that several viruses of importance historically and in modernity such as HIV, rabies, hepatitis B, influenza, and measles all have RNA genomes. The potential value in establishing their evolutionary trajectories, along with the aforementioned *in vivo* processes, makes clear the future utility of ancient RNA.

## Methods

### Samples

To explore the viability of ancient RNA survival, we chose samples considered to have varying potential for success given endogenous DNA content from previous genome analysis [25] but with at least two with a subjectively high potential. Three of the samples represent different tissues (cartilage, liver and muscle) from the same individual: a remarkably well-preserved large canid puppy, with a radiocarbon age of 14,233±34 yBP (ETH-73412; 12,297-12,047 cal BC; 95.4% probability using OxCal v4.2.4 [40], from the village of Tumat in Siberia, Russia. Two puppies were found at the Tumat site, and these analyses concern only puppy #2. (see Table 1). Full descriptions of the samples can be found in Mak et al., 2017 [25]. The three tissue samples from the Tumat puppy were ideal, since they represent varying degrees of preservation from the same individual of advanced ^14^C age. The other two samples, CN214 and CN1921, are both historical skins (hides) from Greenlandic wolves, shot in 1925, and prior to 1869 respectively. Both are currently housed within the Greenland collection at the Natural History Museum of Denmark.

### Laboratory work

All pre-PCR steps of laboratory work including RNA extraction, oligonucleotide processing, and library construction were performed in dedicated ancient DNA facilities equipped with anteroom, and positive air pressure. The ancient DNA facility is physically isolated from PCR areas. All standard approaches to working with ancient biomolecules (PPE clothing, double-layered gloves, deep cleaning, facemasks etc) were followed.

### RNA extraction and purification

Extraction and library construction were performed around protocols designed towards microRNA, due to presumption that it would be necessary to isolate and sequence ultrashort fragments from ancient assemblages given that RNA fragmentation is a time-dependent diagenetic process [11, 15]. RNA was isolated from tissues using an Ambion miRvana kit, following the protocol for total RNA isolation, with the following modifications: prior to digestion, tissues were flash frozen in liquid nitrogen and ground to powder using a mortar and pestle. Tissue powder was then incubated in 1ml of Lysis / Binding buffer for 65 hours at 37°C. Organic extraction with acidic pH 4.2 phenol:chloroform was done to enable phase separation of RNA and DNA [41]. We opted for this method over DNase treatment, because we have previously observed significant inefficiencies of DNase when using ancient DNA as a substrate, often resulting in partial digestion of RNA [42]. We performed organic extraction twice to ensure the purity of RNA, as described [43]. All other steps were performed according to the manufacturer’s instructions; briefly, salt-based precipitation was initiated using a proprietary salt mixture, and consolidated with excess ethanol. RNA was then isolated on a spin-column-attached silica membrane, which was then washed three times using included buffers. RNA was eluted in 50µl, applied at 95°C as per the recommended protocol. The quantity of purified RNA was measured using the Qubit RNA HS assay. Due to known and suspected issues in fluorescence quantification in degraded or fragmented nucleic acid extractions [44], a DNA measurement was not taken using Qubit. We instead opted to measure the level of DNA carryover by quantifying the level of mapping to untranscribed regions of the genome. We subsequently elected to build platform-specific RNA libraries and sequence on two different platforms, the Illumina HiSeq-2500 and the BGISEQ-500, to allow us to explore platform-dependent biases in data generation alongside establishing the survival of ancient RNA.

### Illumina library construction

cDNA libraries were constructed using a NEBNext Multiplex Small RNA Library Prep Set for Illumina according to the manufacturer’s instructions. We opted for this method over other RNA library preparations because of the increased specificity of RNA molecules being incorporated into the library and proven sequence recovery of ultrashort molecules [45]. Briefly, a pre-adenylated 3’ adapter is first ligated to the 5’ end of the RNA molecule. This ATP-free ligation step is facilitated by an RNA ligase mutant, which is truncated to prevent RNA adenylation and thus ligation, unless pre-adenylation of the donor molecule has already occurred [46]. This takes advantage of the 3’ hydroxyl group unique to RNA and thus facilitates enrichment of RNA over potential contaminant DNA. Next, a reverse transcription primer is annealed to the 3’ adapter. Then a standard ssRNA ligation step allows ligation of the 5’ adapter to the RNA molecule to be amplified. Reverse transcription to create single-indexed cDNA libraries based on the RT primer is followed by indexing PCR. Libraries were amplified with between 16 and 20 cycles of PCR using the included polymerase mastermix, and submitted directly for sequencing.

### BGISEQ-500 library construction

For BGISEQ-500 libraries, we utilised the same NEBNext kit with modified adapters and primer oligos appropriate to the BGISEQ-500 platform. We based oligo sequences on those published previously [25] and utilised indexing primers over indexing adapters to reduce costs and improve protocol simplicity, opting for a single 5’ phosphorylated 5’ adapter and adenylated 3’ adapter. Since 5’ adenylation of the 3’ adapter is necessary to RNA-specific library construction as detailed above, the custom BGISEQ-500 3’ adapter was adenylated at the 5’ end using a NEB 5’ Adenylation kit. Libraries were similarly amplified with between 16 and 20 cycles of PCR. With the BGISEQ-500 libraries only, post-PCR products were circularised to form DNB (DNA nanoballs) based on the standard protocol for the platform [25]. DNB production was performed by BGI Europe immediately prior to sequencing.

### Sequencing

Illumina libraries were pooled at equimolar concentrations and sequenced as SE100 on the HiSeq-2500 platform at the Danish National High-Throughput Sequencing Centre. BGI libraries were equally pooled to equimolar concentrations, circularised, and sequenced as SE100 using the BGISEQ-500 platform at BGI Europe, Copenhagen. Demultiplexing was performed in-house and resulting FastQ files were delivered electronically.

### Adapter removal

Illumina and BGI adapters were removed from their respective datasets using cutadapt v.1.11 [47], using default parameters for single-end reads, 10% allowed mismatch, and minimum size retention of 15 nt.

### Read alignment

Sequencing reads from the ancient samples were initially aligned to the CanFam3.1 genome using bowtie2 [48], under default parameters for single-end data. This was done to assess the overall endogenous content including potential DNA contaminants and in relation to previous estimates of endogenous content of the samples [25]. Resulting SAM files were converted to sorted BAM files and filtered by mapping quality score (minimum q=20). The analysis was then repeated using identical parameters, only instead using the CanFam3.1 transcriptome as the reference, and again using canine rRNA and tRNA reference sequences from which to calculate the RNA enrichment factors. Mapping files were de-duplicated, although mapping files with duplicates retained were kept for comparative analyses. Control data was aligned to the CamFam3.1 transcriptome, using default parameters for paired-end data in bowtie2. We performed identical analysis on our extraction blank library and ran any mapped reads through ncbi BLAST+, using default parameters to the nt database, followed with metagenomic analysis using MEGAN to ensure no contamination. All mapped extraction blank reads returned primarily basal or highly conserved assignments, and negligible read numbers were assigned to canids for both Illumina and BGI platforms (2 reads and 39 reads) respectively.

### Junction analysis

We used tophat v2.1.2 [49] to generate an index of exon-exon junctions from the CanFam3.1 genome annotation, and also to map raw, trimmed, de-duplicated RNA-seq reads back to that index. We then collated the frequency of reads straddling exon-exon junctions from the tophat output. We generated intron and exon bedfiles from the CanFam3.1 genome annotation, and used the bedtools intersect function to assess the frequency of reads straddling splice junctions. First, we created a bamfile of reads overlapping exon junctions from our original mapping bamfiles, and fed that output back into bedtools intersect to repeat the analysis, using the intron bedfile instead of the exon bedfile. We used the output from this second round of bedtools intersect to collate read frequencies. We then repeated this analysis using raw, trimmed DNA reads generated previously [25] to compare the two types of data.

### Damage pattern analysis

Cytosine deamination patterns of reads aligned to the CanFam3.1 transcriptome were assessed using mapDamage 2.06 [50]. While the samples had previously showed expected damage patterns from genome sequencing [25], the expectations of similar analysis for RNA are largely unknown due to factors such as single-strandedness and sequence-specific secondary structure formation. We assessed damage profiles on BAM files resulting from both genomic and transcriptomic mapping.

### Control and reference data

For direct transcriptomic comparison, we analysed equivalent, modern NGS data deriving from the same four dog tissue types (skin, cartilage, liver and skeletal muscle). Appropriate data for all tissues was found at the ENA Short Read Archive bioproject accession PRJNA396033, experiment accessions SRX3055179 (cartilage), SRX3055151 (liver), SRX3055143 (skin), and SRX3055142 (muscle). For reference data on relative expression levels between dog tissues, we used Affymetrix array data collated from the Canine Normal Tissue Database, bioproject accession PRJNA124245 [51].

### Expression analysis

Since gene-specific expression analysis has not been performed on ancient material, we attempted two forms of analysis. Method 1 is a direct comparison of control NGS data (see ‘Control and reference data’) to ancient sequencing data. Method 2 was achieved by employing an independent, non-NGS expression array reference [51] to which both modern control NGS and ancient / historical NGS datasets would be compared. Both modern and ancient / historical data was subject to the same analysis.

Both analyses relied on first calculating a relative measure of expression for individual genes within each sample. To generate this, we used the samtools depth function to describe the coverage depth for each position of each transcript, and divided the total coverage for all positions by the length of the transcript to generate a mean coverage value for each. The unique nature of these data creates uncertainties regarding duplicate removal considering excess PCR cycles and short fragments, so we therefore opted to perform analyses using combinations of de-duplicated and duplicates-retained mapping between ancient and control samples. We found that de-duplication, in particular applied to the ancient samples, is more appropriate for these kinds of data (see discussion).

The direct comparison method (method 1) involved firstly performing a variance stabilizing transformation on transcript raw count data, using the Varistran R package (incorporating the edgeR package) [52, 53]. Varistran employs library size normalization using edgeR’s TMM normalization, then applies Anscombe’s [54] variance stabilizing transformation for the negative binomial distribution [52]. Because no replicates were available for each of the ancient samples or controls, dispersion was estimated across the entire dataset (blindly). These normalized data were used for comparison between samples across the entire dataset using Varistran package functions producing ordination biplots and a distance-based heatmap with hierarchal clustering. Biplots were produced by centering rows (genes) by subtracting their global means, performing singular value decomposition and these data plotted where the expression level of a gene in a particular sample, relative to the average expression level of that gene, is approximated by the dot product of the sample position and the gene position (P. Harrison. *Pers. Comm*). Heatmaps were produced by calculating cosine distance, performing hierarchical clustering with *hclust()* and refining clustering using the ‘optimal leaf ordering’ algorithm from the seriation package [55] in order to minimise sharp changes between neighbours without otherwise changing the tree.

To directly compare expression levels between control and ancient/historic samples within and between tissue types, the transformed data for each tissue type were filtered for transcripts within and above the upper 95^th^ percentile of expression levels (i.e. the most highly expressed genes for each tissue type in a given sample). Data below the 95^th^ percentile were discarded, to compensate for noise associated with low-level transcripts [56]. Pairwise linear regression analyses were then performed comparing control tissue expression (explanatory variable) to expression in all ancient /historic tissues (response variable(s)). We corrected for multiple testing [56] using Bonferroni corrections: For each control tissue there were 5 comparisons with ancient / historic samples, so linear models were considered significant at α of 0.01. When comparing control tissues to other control tissues there were 3 comparisons, so linear models were considered significant at α of 0.0166. Linear models between control samples and both ancient and other control samples were only considered relevant if their slope was positive.

For method 2, we first created a simple reference set from the Affymetrix array deriving from the Canine Normal Tissue Database [51]. This was used to describe the tissue to which each annotated gene was most associated with, resulting in a simple gene name to tissue pairing matrix. We then created a second matrix from the CanFam3.1 transcriptome, describing the specific gene name in relation to the gene description (i.e. predicted homology or confirmed). For each sample, we then took transcripts within and above the 95^th^ percentile of expression levels (as calculated earlier using samtools depth) [52, 55, 56] in the sample, we cumulatively scored each of the 10 tissues listed in the Affymetrix array, according to the gene / tissue pairing described in matrix 1. We performed this analysis for all ancient and modern sequencing data, and compared like-for-like sample tissues using a linear regression. We used these analyses to assess the similarity of the modern and ancient datasets based on their appearance when compared to the limited tissue set represented from the Affymetrix array.

### GC content analysis

We assessed the GC content on a per-transcript basis of the CanFam3.1 transcriptome, using a Perl script. We then isolated the transcripts from within the 95^th^ percentile of expression levels as described earlier for consistency. Then, the GC content of individual short reads mapping to those transcripts was calculated on a per-sample basis, from de-duplicated and duplicates-retained bam files (Table S3).

### Metagenomic analysis

For viral infection analysis, we downloaded complete genomes for all available ssRNA and dsRNA viruses known to infect vertebrates from the NCBI Genome resource. Then we mapped all raw reads to the virus dataset using bowtie2, and extracted the mapped reads into fasta format. We then subjected these reads to a full metagenomic BLAST to confirm their viral origin. For tRNA species authentication, we extracted all reads previously mapped to known canine tRNA sequences, and performed a full metagenomic BLAST against the entire nucleotide (nt) database. All BLAST analyses were performed using the NCBI blast+ v.2.6.0 suite, on a standalone high-performance cluster.

## Accession numbers

Control data: Control SRA data for modern transciptomes were taken from the EBI SRA archive, under bioproject PRJNA396033 (see methods).

Our data: All our ancient raw read data was uploaded to the NCBI SRA archive, Accession PRJNA497993.

## Author contributions

OS and MTPG conceived of the study. SF provided the Tumat samples. SF and MG assisted with post-mortem dissections of the Tumat #2 samples. HB provided collagen data and valuable input into molecular preservation theory. MHSS facilitated sample acquisition and valuable input into methods. OS performed all laboratory work, initial NGS data processing, mapping, coverage estimates and all aspects of analytical ‘method 2’. GJD designed and executed all aspects of analytical ‘method 1’. OS, GJD, HV and MTPG wrote the manuscript, with input from all other authors.

## Acknowledgements

The authors wish to thank Professor Robin Allaby at the University of Warwick, for the use of his group’s ancient DNA facility while those at the Centre for GeoGenetics were under renovation, Dr. Roselyn Ware for facilitating laboratory resources and arranging consumables prior to work taking place, and Matthew Poulter/BGI Copenhagen for generating the sequencing data. We also wish to thank Dr. Shyam Gopalakrishnan for his valuable insights on our data analysis and finally we thank Kristian Gregersen at the Natural History Museum of Denmark for access to wolf hides.

---

**Figure S1A:**
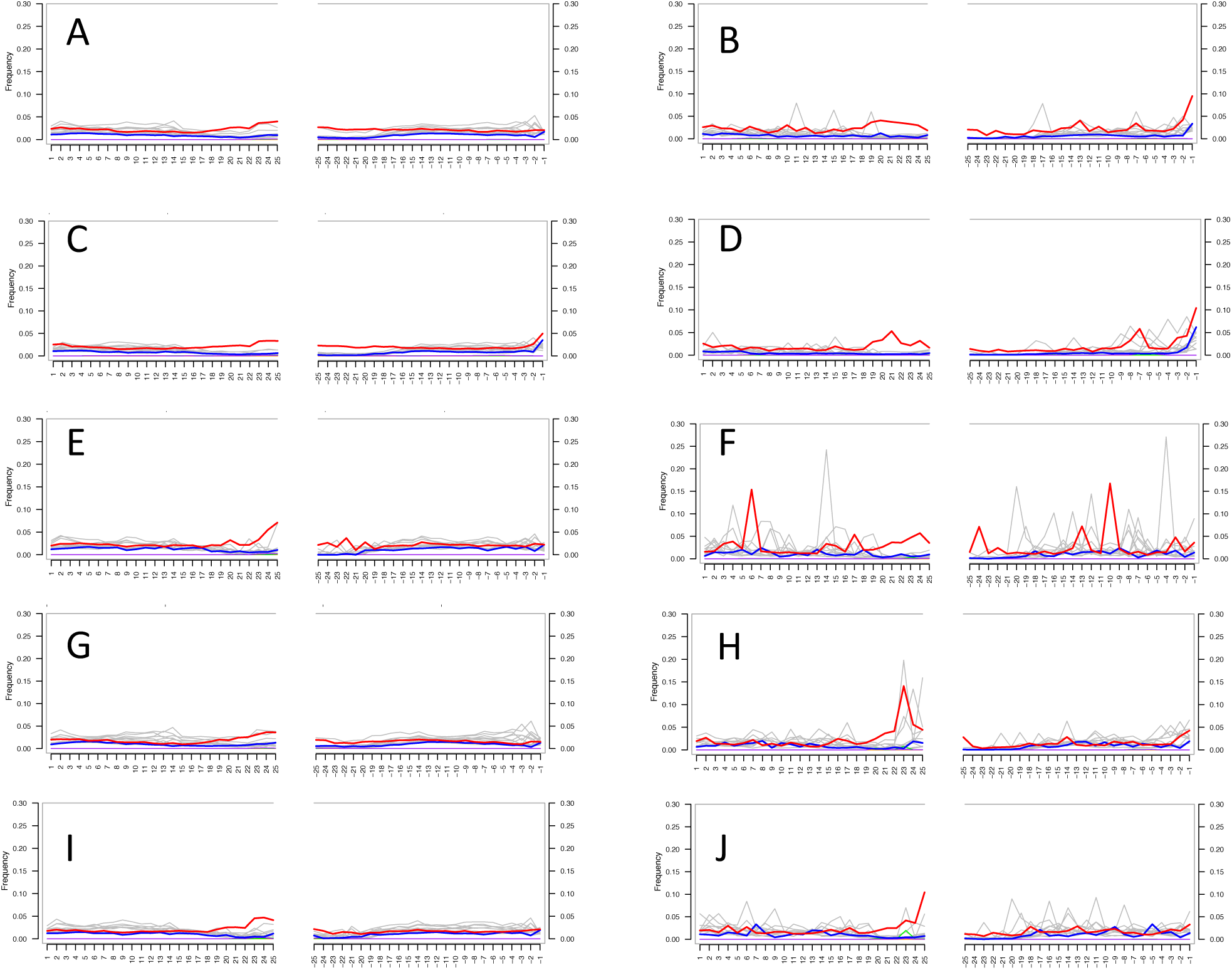
mapDamage profiles of ancient tissues mapped to the CanFam3.1 transcriptome showing nucleotide misincorporations at relative positions from the centre towards the terminal ends of the sequencing read. Red lines indicate C > U misincorporations, blue lines indicate G > A misincorporations, and grey lined indicate others. A) Skin 1, de-duplicated; B) Skin 1, duplicates retained; C) Skin 2, de-duplicated; D) Skin 2, duplicates retained; E) Tumat cartilage, de-duplicated; F) Tumat cartilage, duplicates retained; G) Tumat liver, de-duplicated; H) Tumat liver, duplicated retained; I) Tumat muscle, de-duplicated; J) Tumat muscle, duplicates retained. S1A derived from BGISEG-500 data.

**Figure S1B:**
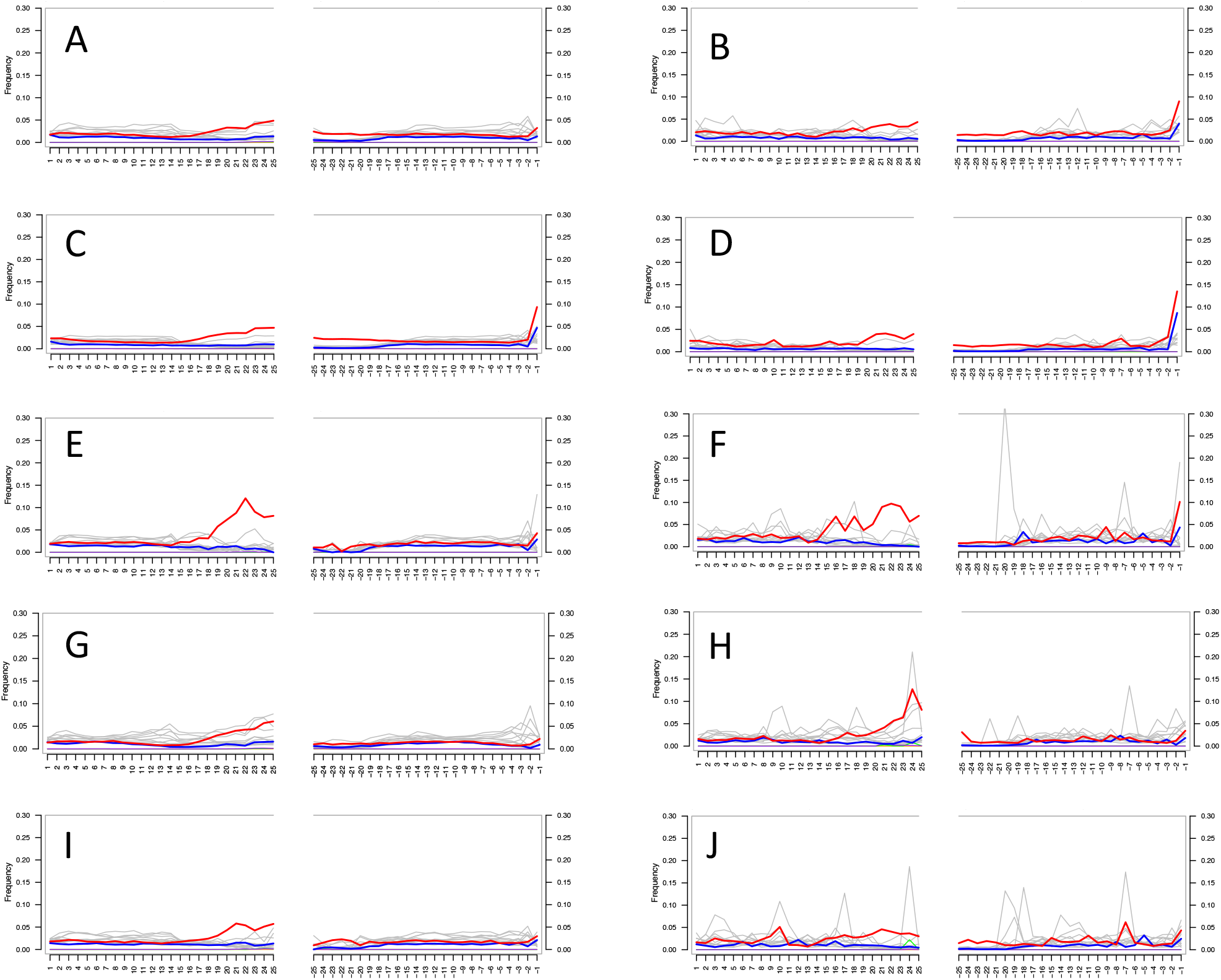
mapDamage profiles of ancient tissues mapped to theCanFam3.1 transcriptome showingnucleotide misincorporations at relative positions from the centre towards the terminal ends of the sequencing read. Redlines indicate C > U misincorporations, blue lines indicate G > A misincorporations,and grey lined indicate others. A) Skin 1, de-duplicated; B) Skin 1, duplicates retained; C) Skin 2, de884 duplicated; D) Skin 2, duplicates retained; E) Tumat cartilage, de-duplicated; F) Tumat cartilage, duplicates retained; G) Tumat liver, de-duplicated; H) Tumat liver, duplicated retained; I) Tumat muscle, de-duplicated; J) Tumat muscle, duplicates retained. S1B from HiSeq-2500 data.

**Figure S2:**
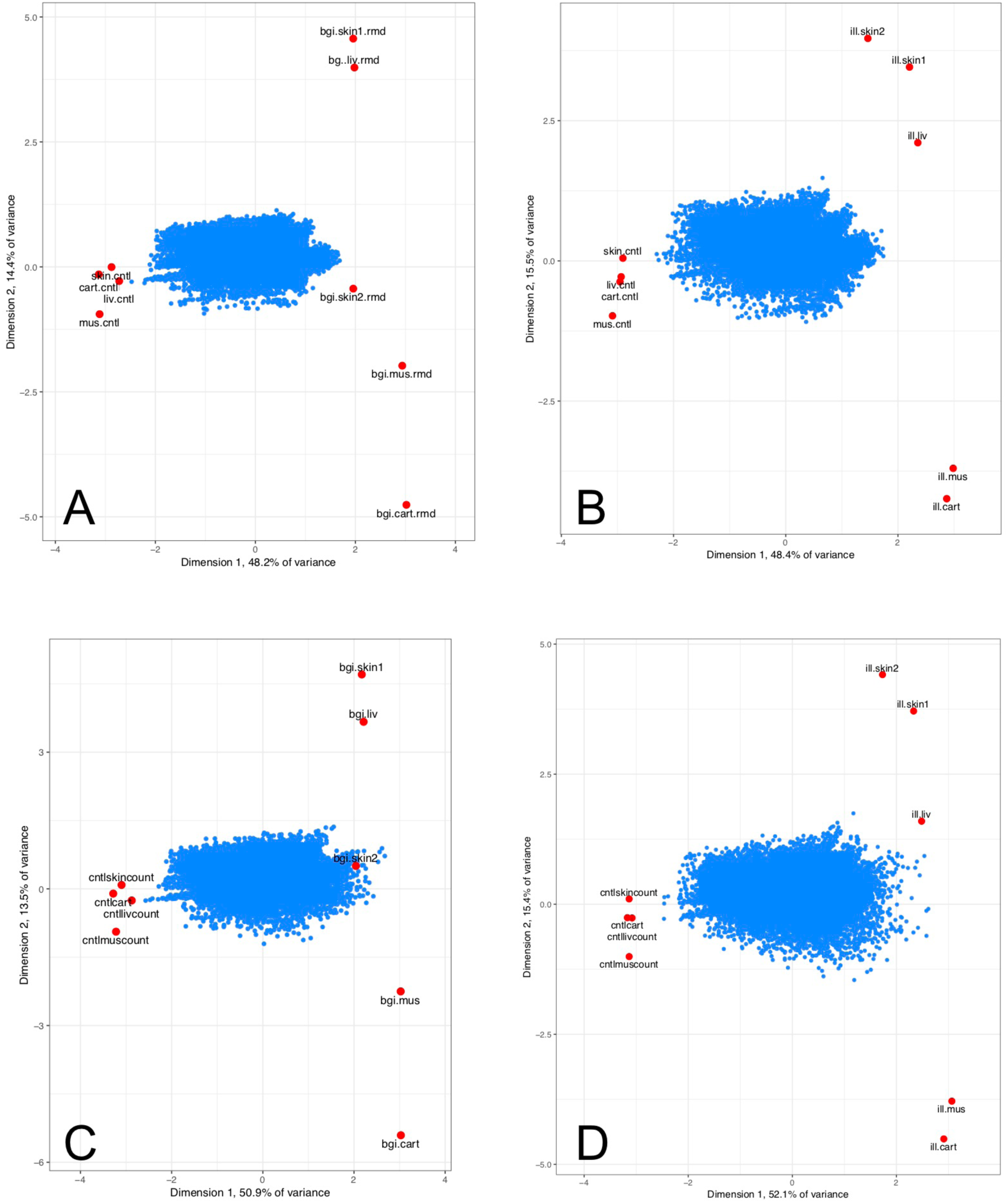
Biplot ordination of standardized individual gene expression (blue points) and similarity between individual samples (red points) along two dimensions (see methods for details). A) BGISEQ-500 data, de-duplicated; B) HiSeq-2500 data, de-duplicated; C) BGISEQ-500 data, duplicates retained; D) HiSeq-2500 data, duplicates retained.

**Figure S3:**
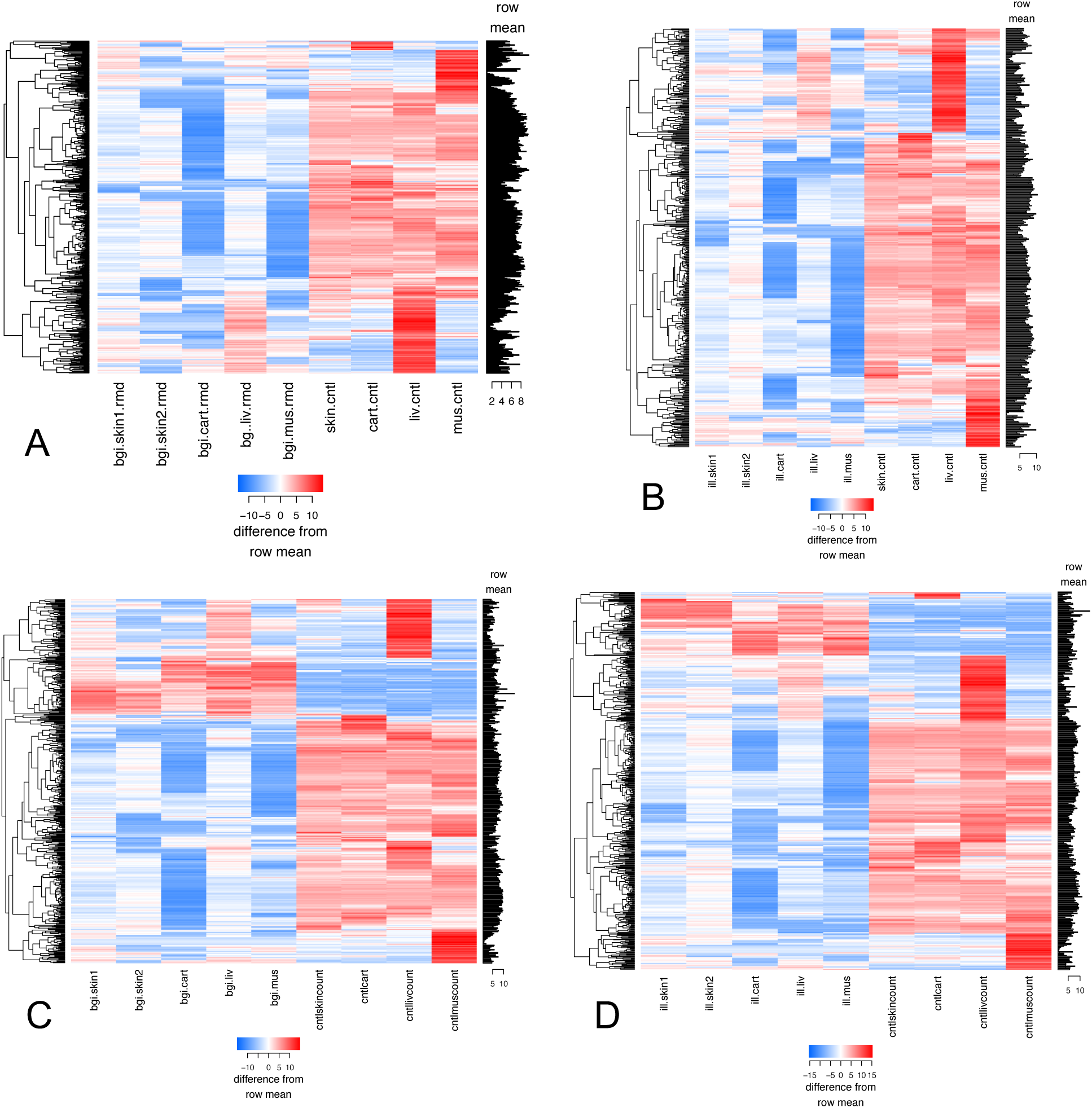
Hierarchical clustering heatmap of similarity between samples (see methods for details) for the top 500 genes with the most differences between samples. A) BGISEQ-500 data, de-duplicated; B) HiSeq-2500 data, de-duplicated; C) BGISEQ-500 data, duplicates retained; D) HiSeq-2500 data, duplicates retained.

**Figure S4:**
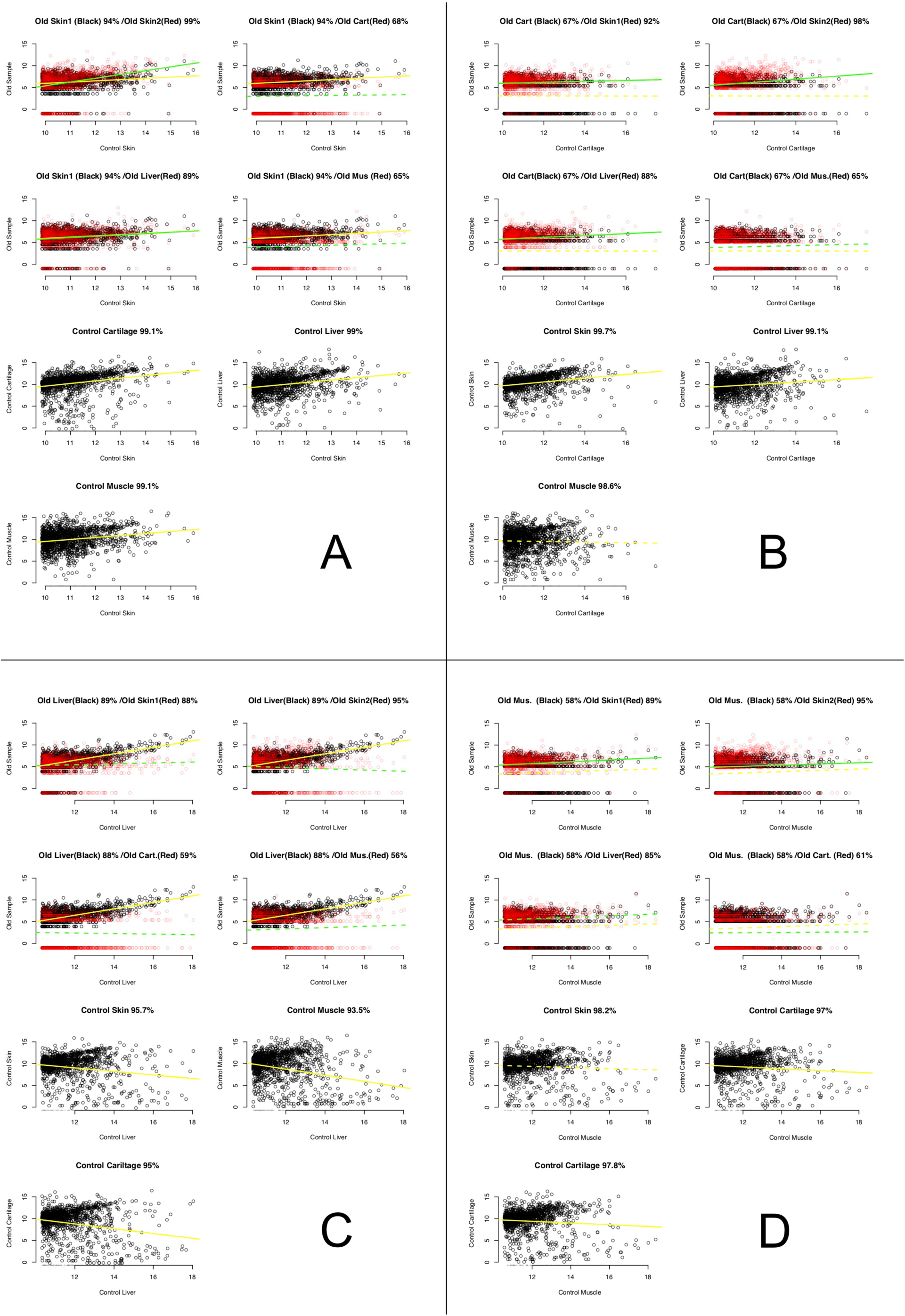
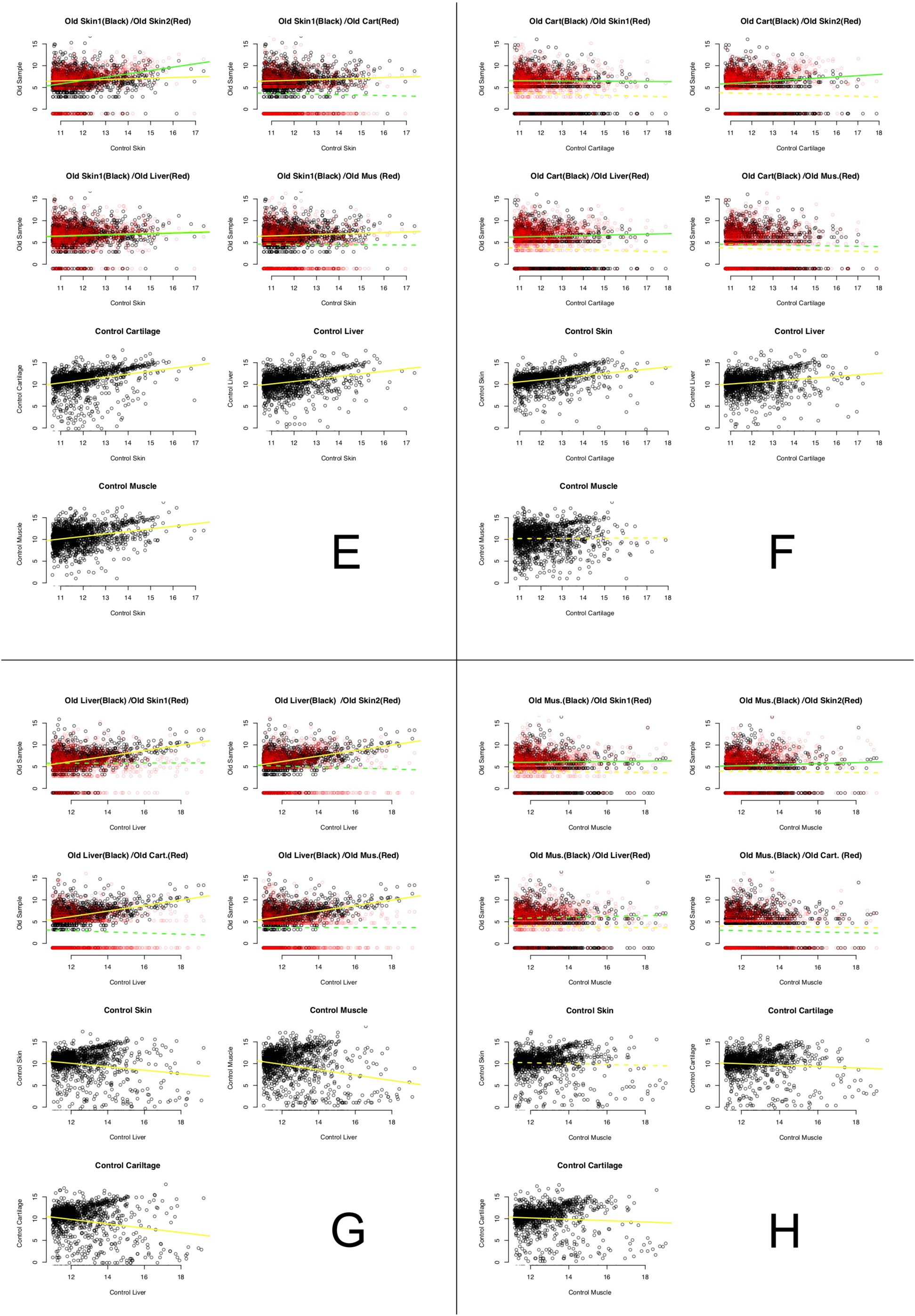
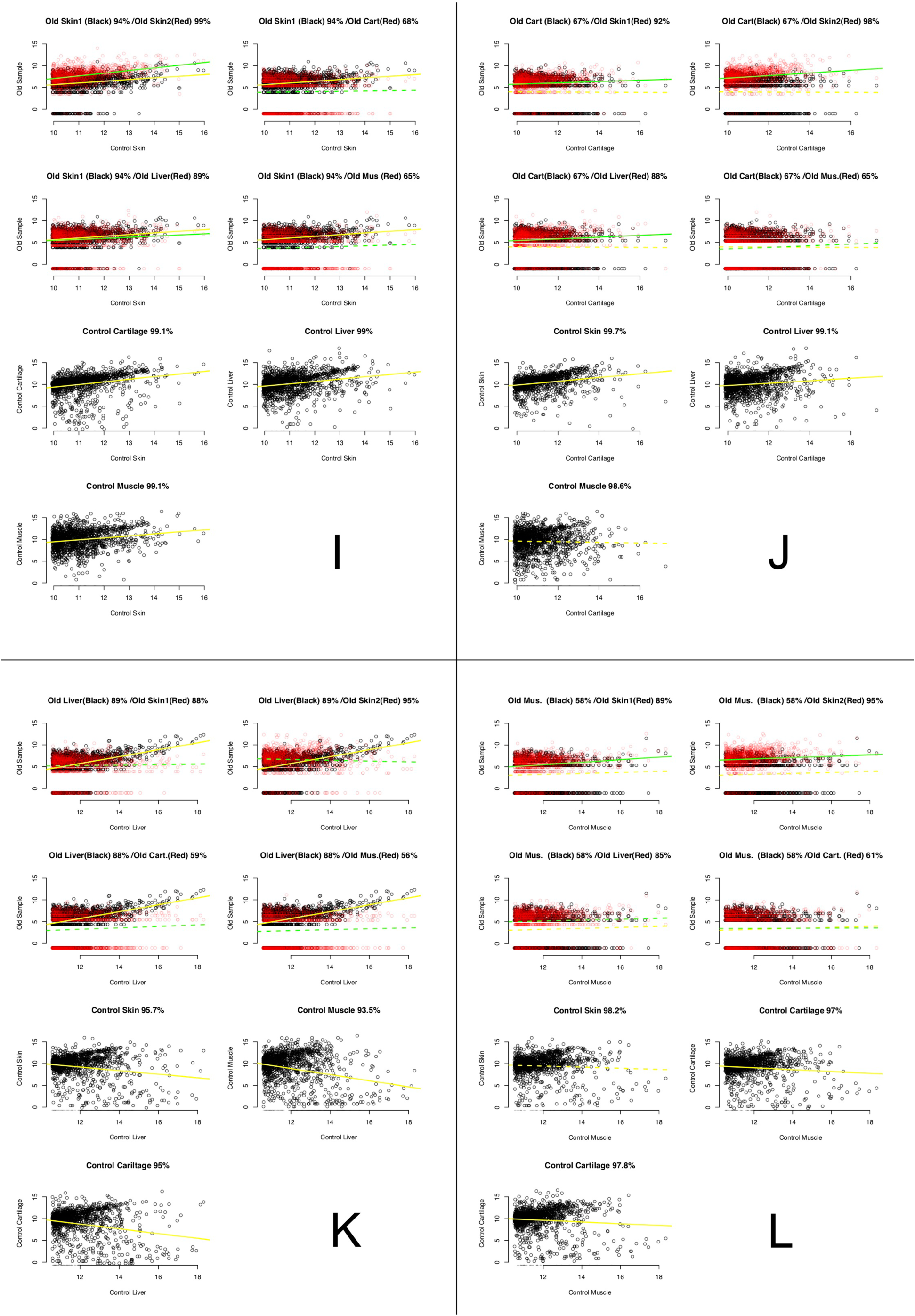
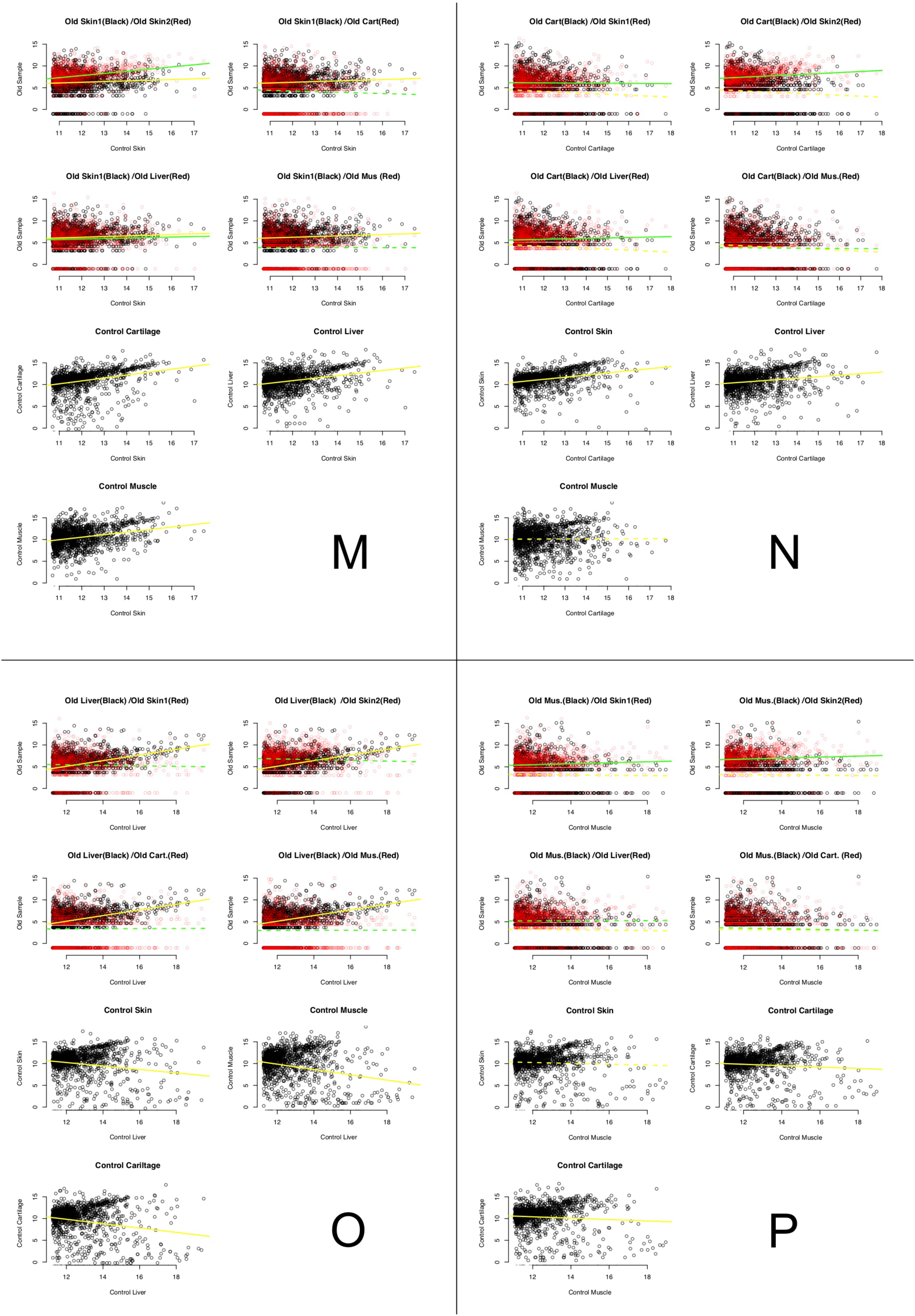
Regressions for all remaining samples, method 1. See legend for Figure 1 for details. A-H, BGISEQ-500; I-P, HiSeq-2500. A-D and I-L, de-duplicated; E-H and M-P, duplicates retained. A, E, I and M, comparison to skin; B, F, J and N, comparison to cartilage; C, G, K and O, comparisons to liver; D, H, L and P, comparisons to muscle.

**Figure S5:**
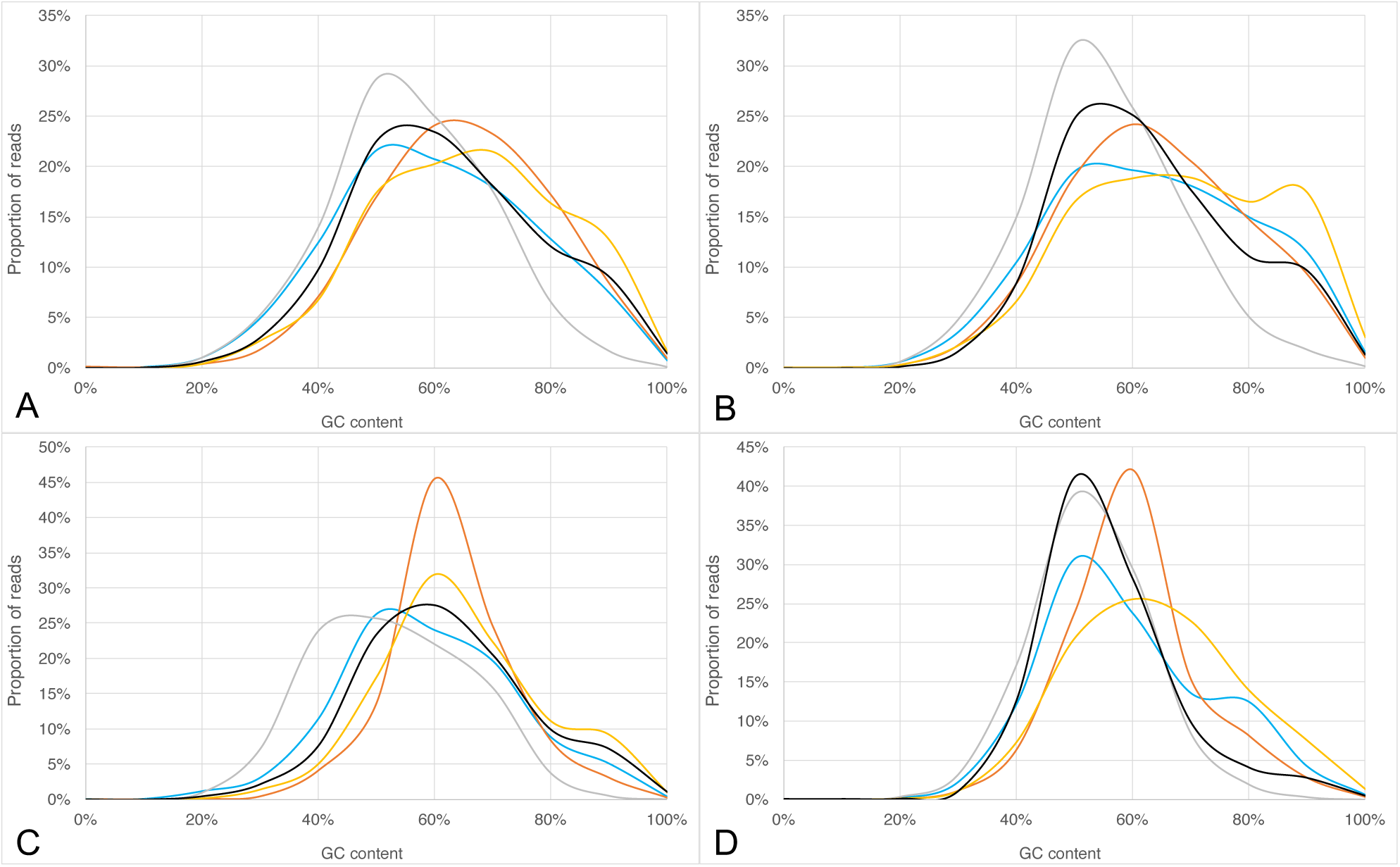
GC content histograms according to sequencing platform and duplicate removal. For all panels: blue line, skin 1; red line, skin 2; grey line, Tumat cartilage; yellow line, Tumat liver; black line, Tumat muscle. A) BGISEQ-500, duplicated removed; B) HiSeq-2500, duplicated removed; C) BGISEQ-500, duplicates retained; D) HiSeq-2500, duplicates retained.

**Figure S6:**
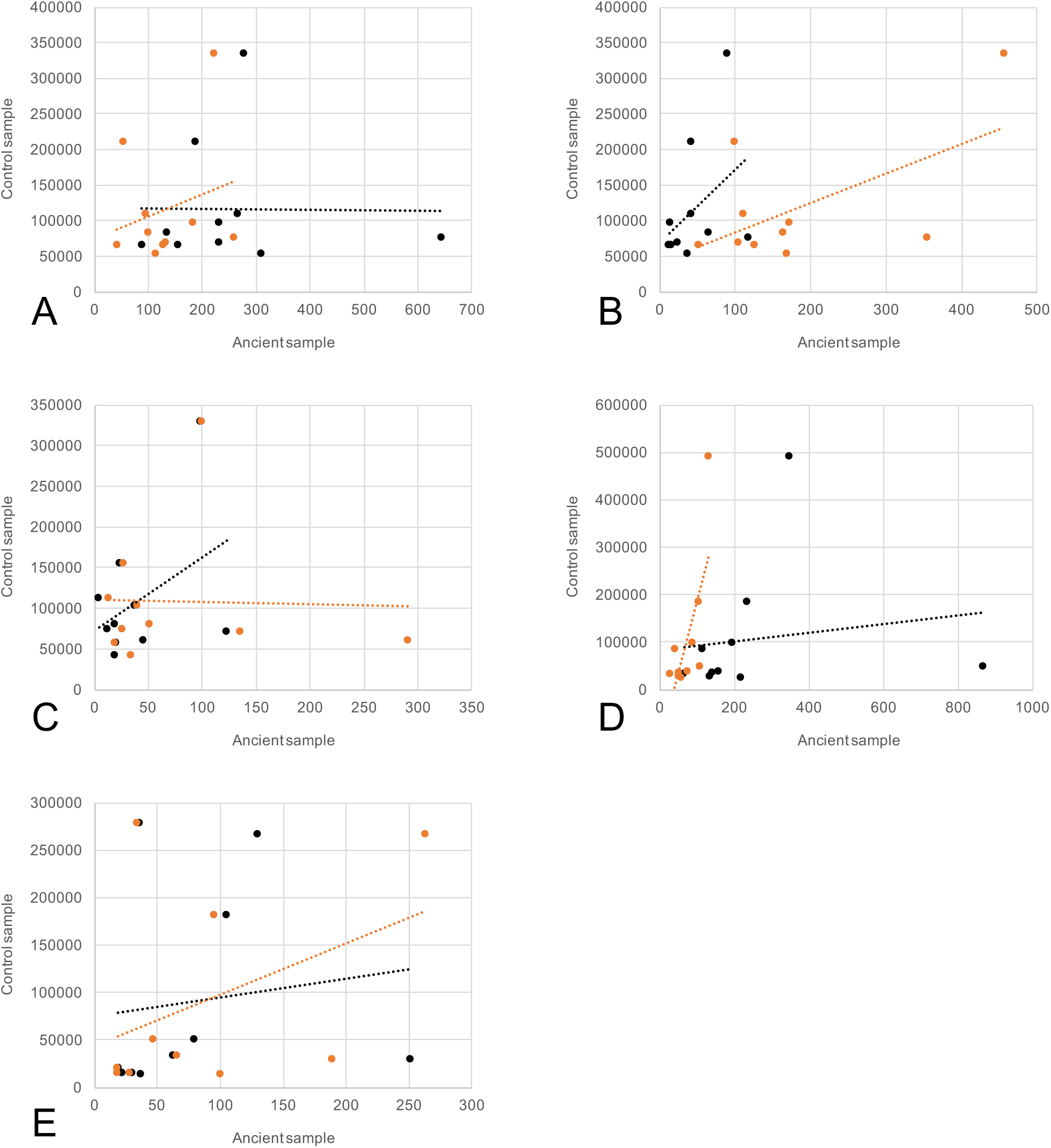
Regressions of all samples, method 2: Relationships between 95% percentile of expressed genes in ancient tissues (x-axis) versus control samples (y-axis). Values are calculated based per-tissue scores (see methods), only retaining duplicate reads. We note here in comparison to duplicate-removed samples that the correlation disintegrates and so suggest for highly amplified libraries, duplicates should be removed. Black data points and trendline refer to BGISEQ-500 data, while orange data points and trendline refer to Illumina HiSeq-2500 data. A) Skin 1; B) Skin 2; C) Tumat cartilage; D) Tumat liver; E) Tumat muscle

**Figure S7:**
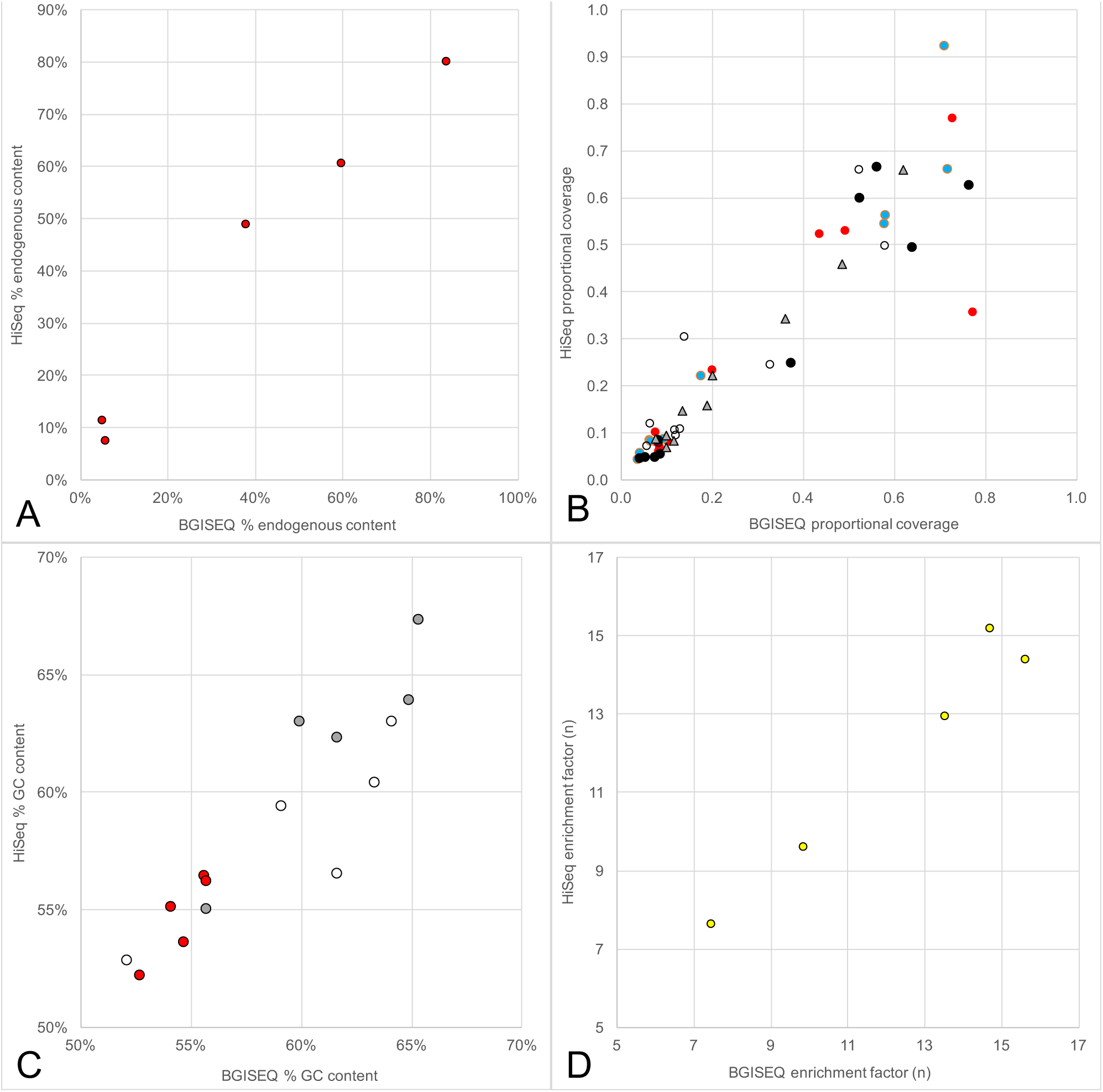
comparison of data generated by BGISEQ-500 and HiSeq-2500 platforms. A) endogenous content of sequencing reads by tissue (see Table S2). B) Regressions of method 2 between platforms. Red circles, Skin 1; white circles, Tumat cartilage; blue circles, Skin 2; black circles, Tumat liver; grey triangles, Tumat muscle. C) Mean GC content of reads by tissue, depending on duplication. Red circles, reads mapping to the 95^th^ percentile and above of expression after mapping and deduplication. White circles, all mapped reads with deduplication. Grey circles, all mapped reads without deduplication. D) RNA enrichment factor by tissue type.

**Figure S8A:**
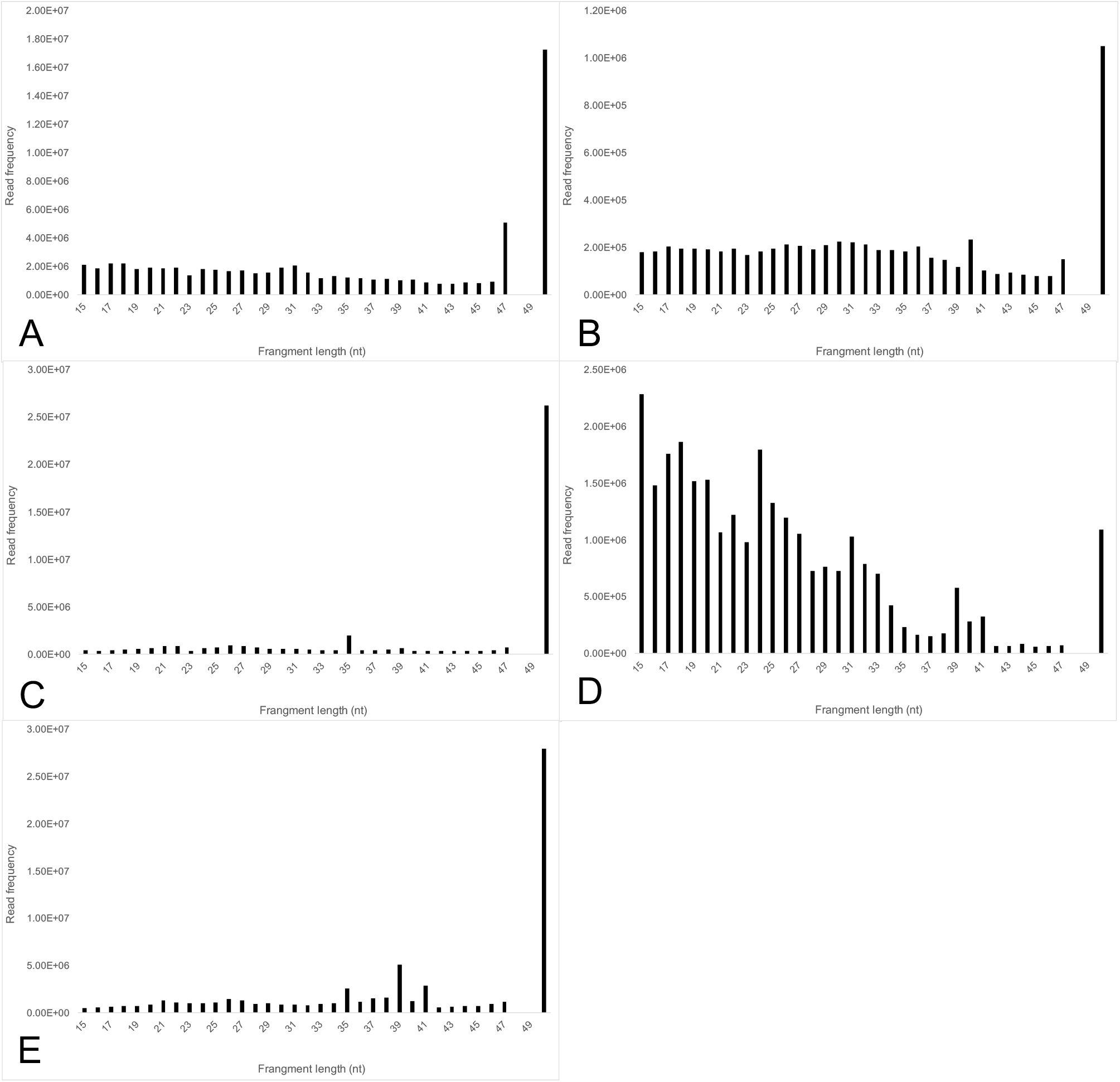
length distribution plots of BGISEQ-500 RNA-seq. A) Skin 1; B) Skin 2; C) Tumat cartilage; D) Tumat liver; E) Tumat muscle.

**Figure S8A:**
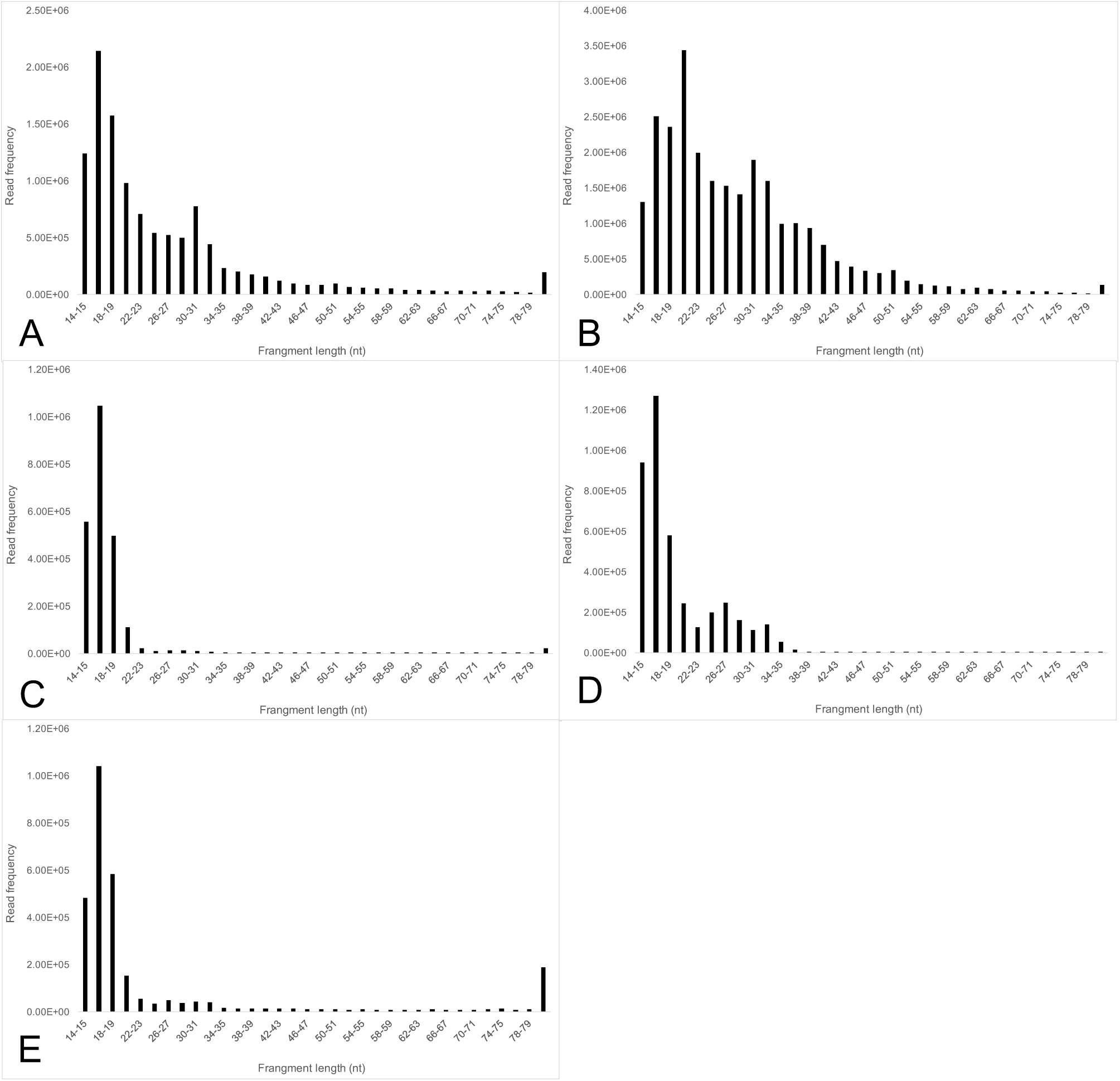
length distribution plots of HiSeq-2500 RNA-seq. A) Skin 1; B) Skin 2; C) Tumat cartilage; D) Tumat liver; E) Tumat muscle.

**Figure S9A:**
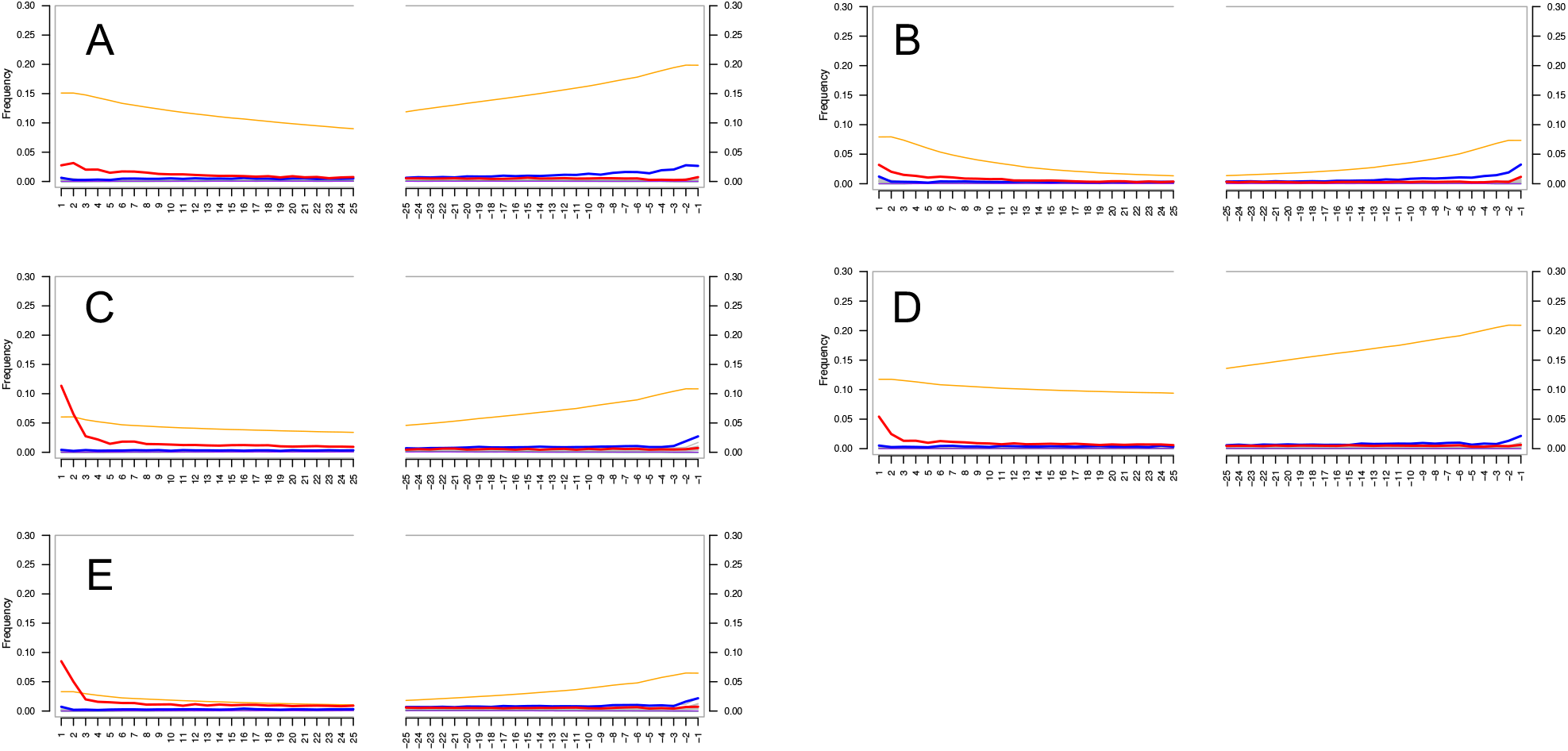
mapDamage plots of DNA data from Mak et al 2018 sequenced on the BGISEQ-500 plaform. A) Skin 1; B) Skin 2; C) Tumat cartilage; D) Tumat liver; E) Tumat muscle.

**Figure S9A:**
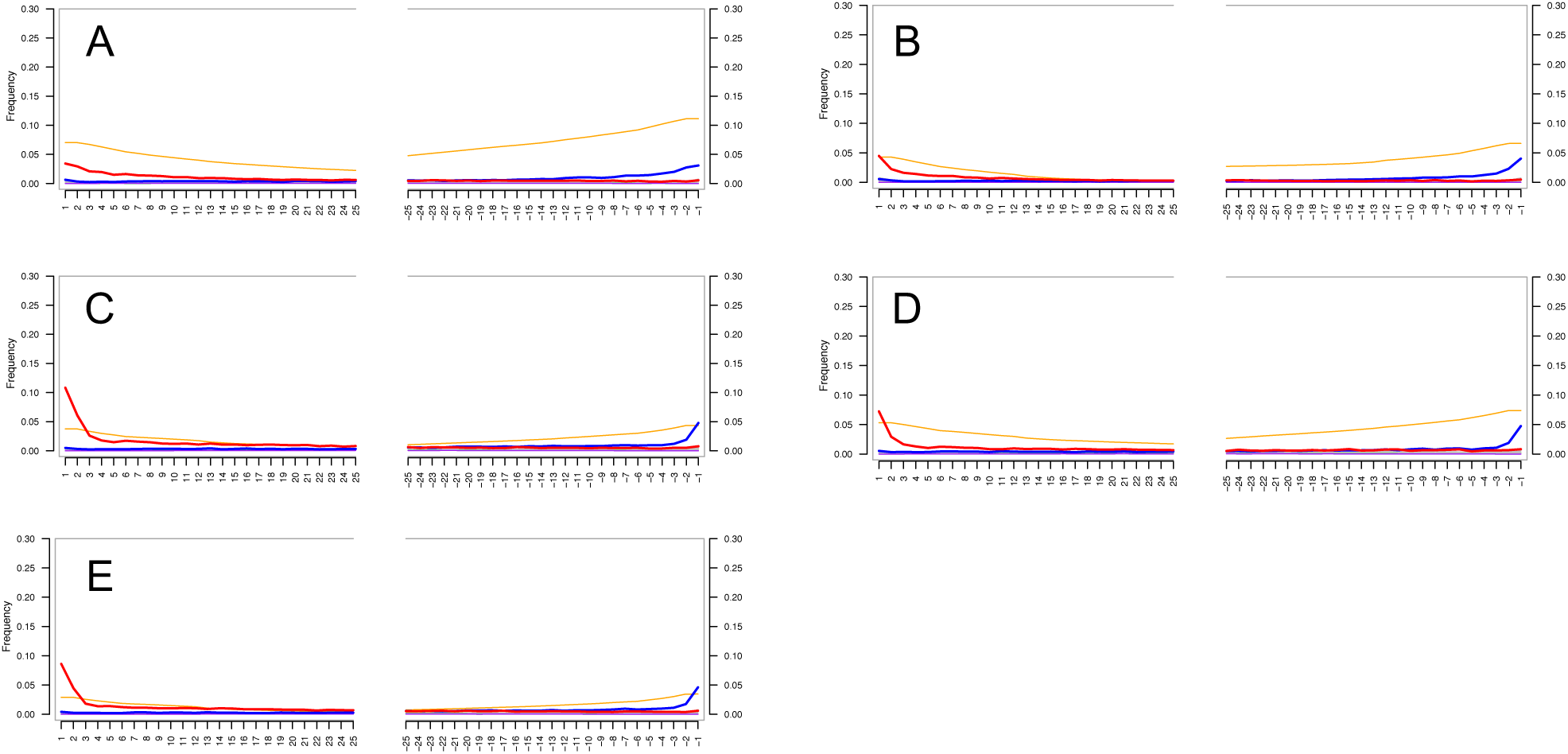
mapDamage plots of DNA data from Mak et al 2018 sequenced on the HiSeq-2500 plaform. A) Skin 1; B) Skin 2; C) Tumat cartilage; D) Tumat liver; E) Tumat muscle.

**Table S1:** Junction analysis of RNA-seq and DNA data derived from the same samples. Reads mapping over splice junctions and exon-exon junctions were collated for each sample and molecule type, and enrichment factors calculated. In all cases, RNA-seq data shows significantly more exon-exon junction coverage than splice junctions, highlighting it’s authenticity. Conversely, the opposite trend is seen for DNA data.

| Sample | RNA |  |  | DNA |  |  |
| --- | --- | --- | --- | --- | --- | --- |
|  | Splice junction | Exon/Exon | Enrichment factor | Splice junction | Exon/Exon | Enrichment factor |
| Skin 1 BGI | 2,560 | 219,511 | 85.75 | 239,562 | 169,698 | 0.71 |
| Skin 2 BGI | 1,491 | 158,582 | 106.36 | 12,765,554 | 369,114 | 0.03 |
| Tumat cartilage BGI | 498 | 1,831 | 3.68 | 588,823 | 14,259 | 0.02 |
| Tumat liver BGI | 2,164 | 270,239 | 124.88 | 24,981 | 422 | 0.02 |
| Tumat muscle BGI | 969 | 4,289 | 4.43 | 1,841,006 | 49,364 | 0.03 |
| Skin 1 HiSeq | 1,765 | 112,064 | 63.49 | 172,280 | 90,753 | 0.53 |
| Skin 2 HiSeq | 4,066 | 756,268 | 186.00 | 91,479 | 20,184 | 0.22 |
| Tumat cartilage HiSeq | 641 | 3,405 | 5.31 | 277,720 | 1,768 | 0.01 |
| Tumat liver HiSeq | 1,495 | 102,469 | 68.54 | 3,069 | 476 | 0.16 |
| Tumat muscle HiSeq | 786 | 7,304 | 9.29 | 508,984 | 27,548 | 0.05 |

**Table S2:** Method 2 final scores according to Affymetrix array tissue derived from modern and ancient NGS datasets. Top half, scores following deduplication. Lower half, scores with duplicate reads retained.

|  |  | Tissue |  |  |  |  |  |  |  |  |  |
| --- | --- | --- | --- | --- | --- | --- | --- | --- | --- | --- | --- |
| Sample, duplicates removed |  | Brain | Heart | Jejunum | Kidney | Liver | Lung | Lymphnode | Pancreas | Skel muscle | Spleen |
|  | Skin 1 BGI | 7.369512 | 9.224989 | 5.976252 | 6.06709 | 5.788138 | 12.711885 | 22.262183 | 14.300985 | 10.964448 | 3.526031 |
|  | Skin 2 BGI | 4.703452 | 6.649602 | 6.497142 | 8.674943 | 3.252891 | 14.391111 | 48.378053 | 11.058813 | 4.697513 | 2.944075 |
|  | Tumat cartilage BGI | 0.6524631 | 0.4191422 | 0.8122769 | 0.7963115 | 0.347326 | 0.6237626 | 1.5482203 | 1.8898358 | 0.7800282 | 0.1445928 |
|  | Tumat liver BGI | 5.867459 | 10.052321 | 6.673093 | 9.534536 | 56.858722 | 4.42351 | 17.627531 | 16.918639 | 8.540282 | 3.965165 |
|  | Tumat muscle BGI | 1.4682131 | 1.008993 | 1.2243416 | 1.5066447 | 1.1267399 | 1.6271386 | 1.4323754 | 2.8535713 | 1.9912942 | 0.5152724 |
|  | Skin 1 HiSeq | 4.502981 | 3.257765 | 2.945736 | 2.025289 | 2.246489 | 7.007347 | 11.993489 | 5.869402 | 4.173192 | 0.633882 |
|  | Skin 2 HiSeq | 20.837246 | 29.993212 | 26.340306 | 26.351402 | 11.292727 | 59.382402 | 140.366709 | 40.869815 | 17.477253 | 14.142799 |
|  | Tumat cartilage HiSeq | 0.9120192 | 0.7508219 | 1.0398841 | 0.9746833 | 0.9679512 | 0.684925 | 1.6248867 | 2.574598 | 1.7789093 | 0.3543757 |
|  | Tumat liver HiSeq | 2.952995 | 3.015451 | 2.740994 | 4.73437 | 31.771138 | 1.163108 | 5.040153 | 10.595909 | 2.663559 | 1.860064 |
|  | Tumat muscle HiSeq | 1.2044026 | 1.01611 | 0.7344993 | 1.4796762 | 0.7128224 | 1.2645689 | 1.5379421 | 2.5229572 | 2.0718783 | 0.4429613 |
|  | skin_ctrl | 45415.554 | 50561.467 | 30482.011 | 40899.5 | 26136.132 | 104940.71 | 167978.585 | 40188.386 | 35487.522 | 28793.465 |
|  | liver_ctrl | 30122.055 | 59834.033 | 27707.201 | 79697.661 | 374555.123 | 20205.328 | 107564.365 | 38311.08 | 18638.103 | 22270.378 |
|  | muscle_ctrl | 41331.829 | 203341.122 | 12626.31 | 27125.846 | 11126.786 | 12401.362 | 108659.232 | 24176.597 | 210645.542 | 13996.461 |
|  | cart_ctrl | 72084.93 | 48243.49 | 36999.322 | 53199.036 | 25481.206 | 101362.97 | 188636.671 | 47892.239 | 42075.641 | 72851.855 |
| Sample, duplicates retained |  | Brain | Heart | Jejunum | Kidney | Liver | Lung | Lymphnode | Pancreas | Skel muscle | Spleen |
|  | Skin 1 BGI | 230.804808 | 265.141447 | 155.092406 | 133.99913 | 308.581048 | 187.204974 | 278.045287 | 643.430022 | 231.321085 | 87.978841 |
|  | Skin 2 BGI | 13.408641 | 41.460159 | 12.134565 | 64.229466 | 36.629826 | 42.227093 | 89.674146 | 116.653709 | 23.490457 | 14.965661 |
|  | Tumat cartilage BGI | 37.445289 | 11.365473 | 19.945933 | 19.373542 | 18.559948 | 23.404146 | 98.726107 | 123.02492 | 45.598856 | 4.045317 |
|  | Tumat liver BGI | 157.97178 | 112.81358 | 140.34369 | 193.27382 | 346.1022 | 134.3815 | 232.51709 | 867.15393 | 216.46818 | 64.93643 |
|  | Tumat muscle BGI | 79.413814 | 35.918867 | 30.175458 | 62.279461 | 36.953227 | 22.080463 | 104.534059 | 250.856177 | 129.572004 | 18.72993 |
|  | Skin 1 HiSeq | 182.49211 | 95.13863 | 127.08658 | 100.47759 | 112.55379 | 53.95795 | 222.97553 | 259.54267 | 131.5064 | 41.83447 |
|  | Skin 2 HiSeq | 171.95332 | 111.24421 | 126.00464 | 163.85154 | 168.81874 | 99.18516 | 455.93314 | 354.99892 | 104.96765 | 50.92724 |
|  | Tumat cartilage HiSeq | 39.717703 | 25.174315 | 19.145569 | 51.30038 | 33.601261 | 27.438784 | 99.677509 | 135.633345 | 291.15729 | 13.595632 |
|  | Tumat liver HiSeq | 73.421856 | 42.600503 | 49.515499 | 88.649362 | 131.68405 | 52.394247 | 103.138827 | 108.894429 | 57.308926 | 27.586459 |
|  | Tumat muscle HiSeq | 46.617127 | 34.137448 | 27.737057 | 65.530679 | 99.727602 | 18.514318 | 95.543809 | 188.81023 | 263.027508 | 18.130081 |
|  | skin_ctrl | 97570.01 | 109034.171 | 66239.344 | 83640.604 | 53400.6 | 210011.027 | 333949.46 | 75736.27 | 69766.985 | 65994.027 |
|  | liver_ctrl | 37598.592 | 85657.563 | 36832.959 | 99082.865 | 490673.756 | 27588.329 | 184312.883 | 49950.749 | 24389.728 | 32640.816 |
|  | muscle_ctrl | 50403.683 | 278563.469 | 15797.42 | 34023.955 | 13444.188 | 15634.505 | 180908.049 | 29621.29 | 267055.102 | 19846.433 |
|  | cart_ctrl | 103660.907 | 74625.811 | 57230.311 | 80891.516 | 41995.311 | 155869.564 | 329032.457 | 71605.872 | 59867.114 | 112521.816 |

**Table S3:** Mean GC content of mapped reads depending on selection and (de)duplication.

| Sample | 95 %ile GC | Overall Read GC,<br>duplicates removed | Overall Read GC,<br>duplicates retained |
| --- | --- | --- | --- |
| Skin 1 BGI | 54.1 | 59.1 | 59.9 |
| Skin 2 BGI | 55.6 | 63.3 | 64.9 |
| Tumat cartilage BGI | 52.7 | 52.1 | 55.7 |
| Tumat liver BGI | 55.7 | 64.1 | 65.3 |
| Tumat muscle BGI | 54.7 | 61.6 | 61.6 |
| Skin 1 HiSeq | 55.1 | 59.4 | 63 |
| Skin 2 HiSeq | 56.4 | 60.4 | 63.9 |
| Tumat cartilage HiSeq | 52.2 | 52.8 | 55 |
| Tumat liver HiSeq | 56.2 | 63 | 67.3 |
| Tumat muscle HiSeq | 53.6 | 56.5 | 62.3 |

**Table S4:** Basic NGS statistics of DNA data, subjected to the same analysis as the RNA-seq data of the same samples. Note that the ribosomal RNA proportion and overall RNA enrichment factors are significantly less than those of the RNA-seq data.

|  | Sample # | Species | Tissue | Age | Genome | mRNA | rRNA | Proportion<br>rRNA | tRNA | RNA Enrichment<br>factor |
| --- | --- | --- | --- | --- | --- | --- | --- | --- | --- | --- |
| BGISEQ | Skin 1 | Wolf | Skin | Before 1869 AD | 88,606,127 | 3,400,335 | 138,318 | 0.15% | 198,399 | 0.58 |
|  | Skin 2 | Wolf | Skin | 1925 AD | 19,539,088 | 1,499,806 | 34,885 | 0.16% | 183,823 | 1.21 |
|  | Tumat C | Canid | Cartilage | ca. 14122 YBP | 28,894,255 | 486,848 | 19,637 | 0.07% | 939 | 0.24 |
|  | Tumat L | Canid | Liver | ca. 14122 YBP | 1,252,563 | 37,439 | 1,934 | 0.15% | 674 | 0.44 |
|  | Tumat M | Canid | Muscle | ca. 14122 YBP | 89,229,030 | 1,504,208 | 61,956 | 0.07% | 3,125 | 0.24 |
| HiSeq | Skin 1 | Wolf | Skin | Before 1869 AD | 7,006,239 | 304,201 | 12,334 | 0.17% | 25,443 | 0.67 |
|  | Skin 2 | Wolf | Skin | 1925 AD | 14,216,858 | 966,092 | 26,558 | 0.17% | 143,140 | 1.10 |
|  | Tumat C | Canid | Cartilage | ca. 14122 YBP | 1,622,174 | 34,365 | 1,552 | 0.09% | 208 | 0.31 |
|  | Tumat L | Canid | Liver | ca. 14122 YBP | 201,084 | 7,820 | 285 | 0.14% | 203 | 0.57 |
|  | Tumat M | Canid | Muscle | ca. 14122 YBP | 29,592,985 | 632,765 | 30,098 | 0.10% | 4,750 | 0.31 |

Supplementary Data 1 (see supplementary data excel file Supp_Data_1.xlsx): Regression table of Method 1. Details of linear regression analysis of the 95th percentile of genes expressed in each control tissue, compared with each ancient tissue and other control tissues. Models marked in bold have the slope in the expected direction (positive) and are significant at bonferroni alphas adjusted for multiple comparisons (ancient tissues alpha = 0.01, control tissues alpha = 0.0166).

Supplementary Data 2 (see supplementary data files Supp_Data_2_dupsRemoved.xlsx and Supp_Data_2_dupsRetained.xlsx on Google Drive at https://drive.google.com/open?id=1cO88r8RrjLRGOnA80hdy6TGVH-eUppH4): Scoring matrix for method 2 arranged in tabs by tissue and sequencing platform. Briefly: columns A and B are the static tissue/gene pairs generated from the Canine Normal Tissue Database (CNTD) Affymetrix array. Column D is the NCBI reference for each gene found on the CanFam3.1 transcriptome, column F the full gene description, and column G the derived gene name / loc ID. Column E is the mean coverage depth of that gene after mapping. Column H is a lookup formula to assign each gene a most-related tissue from the 10 listed on CNTD. Column I is the 95^th^ percentile value of coverage. Columns J-S are the total cumulative scores assigned to each of the 10 tissues following associated-gene / score pairing. One data file is for analysis with de-duplicated data (dupsRemoved), the other with duplicates retained (dupsRetained).

